# Modeling Cortical Versus Hippocampal Network Dysfunction in a Human Brain Assembloid Model of Epilepsy and Intellectual Disability

**DOI:** 10.1101/2024.09.07.611739

**Authors:** Colin M. McCrimmon, Daniel Toker, Marie Pahos, Kevin Lozano, Jack J. Lin, Jack Parent, Andrew Tidball, Jie Zheng, László Molnár, Istvan Mody, Bennett G. Novitch, Ranmal A. Samarasinghe

**Affiliations:** Department of Neurology, University of California, Los Angeles, David Geffen School of Medicine, Los Angeles, 90095, CA, USA; Department of Neurology, University of California, Davis, School of Medicine, Sacramento, 95817, CA, USA; Center for Mind and Brain, University of California, Davis, Davis, 95618, CA, USA; Department of Neurology, University of Michigan School of Medicine, Ann Arbor, 48109, MI, USA; Michigan Neuroscience Institute, University of Michigan, Ann Arbor, 48109, MI, USA; VA Ann Arbor Healthcare System, Ann Arbor, 48105, MI, USA; Department of Biomedical Engineering, University of California, Davis, Davis, 95616, CA, USA; Department of Neurological Surgery, University of California, Davis, School of Medicine, Sacramento, 95817, CA, USA; Department of Electrical Engineering, Sapientia Hungarian University of Transylvania, Târgu-Mureş/Corunca, 540485, ROU; Department of Physiology, University of California, Los Angeles, David Geffen School of Medicine, Los Angeles, 90095, CA, USA; Department of Neurobiology, University of California, Los Angeles, Los Angeles, 90095, CA, USA; Eli and Edythe Broad Center for Regenerative Medicine and Stem Cell Research, University of California, Los Angeles, Los Angeles, 90095, CA, USA; Intellectual Development and Disabilities Research Center, University of California, Los Angeles, Los Angeles, 90095, CA, USA

## Abstract

Neurodevelopmental disorders often impair multiple cognitive domains. For instance, a genetic epilepsy syndrome might cause seizures due to cortical hyperexcitability and present with memory impairments arising from hippocampal dysfunction. This study examines how a single disorder differentially affects distinct brain regions by using human patient iPSC-derived cortical- and hippocampal-ganglionic eminence assembloids to model Developmental and Epileptic Encephalopathy 13 (DEE-13), a condition arising from gain-of-function mutations in the *SCN8A* gene. While cortical assembloids showed network hyperexcitability akin to epileptogenic tissue, hippocampal assembloids did not, and instead displayed network dysregulation patterns similar to in vivo hippocampal recordings from epilepsy patients. Predictive computational modeling, immunohistochemistry, and single-nucleus RNA sequencing revealed changes in excitatory and inhibitory neuron organization that were specific to hippocampal assembloids. These findings highlight the unique impacts of a single pathogenic variant across brain regions and establish hippocampal assembloids as a platform for studying neurodevelopmental disorders.

## 1 Introduction

Brain organoids derived from human induced pluripotent stem cells (hiPSCs) and human embryonic stem cells (hESCs) are increasingly used as models of complex neurological disorders [1–3]. Historically, brain organoid studies focused on structural, cellular, and transcriptomic consequences of neural pathology. More recent studies have highlighted the ability of organoids to recapitulate key features of network-wide neural electrodynamics, like oscillatory field potentials with spectral peaks at frequencies seen in the human brain in vivo [4–6]. Importantly, organoid models of neurological conditions with single gene pathogenic variants can recreate aberrant neural electrodynamics [4]. This indicates that organoids can be used to study the impact of neurological disease on circuit composition and network-level electrical activity — features central to disease pathology that are difficult to study in vivo. The existence of robust protocols for generating brain-region specific organoids [7–11] further enables detailed exploration of the impact of genetic perturbations on brain network development and function in distinct regions. Given the complex phenotypes and multiple comorbidities of many neurological disorders, this may allow for a better understanding of disease pathophysiology and more targeted management of comorbidities.

We demonstrate the value of this approach in understanding network development and function in Developmental and Epileptic Encephalopathy 13 (DEE-13), a condition arising from gain-of-function mutations in the *SCN8A* gene (encoding sodium channel Nav1.6). DEE-13, like all DEEs, typically features early-onset epilepsy and intellectual disability [12, 13]. These symptoms are hypothesized to stem from independent developmental perturbations, though the mechanisms underlying epilepsy on the one hand and intellectual disability on the other remain unclear [12]. To understand how *p*.*SCN8A* variants impact network development in different brain regions, we generated an organoid model of cortex and hippocampus using two *p*.*SCN8A* variants and an isogenic control. To create structures with integrated excitatory and inhibitory neurons and glia, essential for complex electrodynamic network activity [4], we generated Cx+GE assembloids using established protocols [4, 10] and also developed hippocampus-ganglionic eminence (Hc+GE) assembloids.

Using local field potentials and calcium imaging, we found overt hyperexcitability in Cx+GE, consistent with prior DEE-13 mouse models [14] and hiPSC-derived 2D neuronal models [15]. Surprisingly, Hc+GE assembloids did not show overt hyperexcitability, but instead exhibited aberrations of network activity patterns that are associated with normal hippocampal function. Specifically, we observed disordered theta-gamma phase-amplitude coupling, reduced high gamma/ripple frequency oscillations, and destabilized dynamics at single-cell and circuit levels in Hc+GE assembloids with gain-of-function *p*.*SCN8A* variants. Analysis of in vivo human hippocampal oscillations from temporal lobe epilepsy patients revealed similar aberrations in phaseamplitude coupling in epileptic hippocampi compared to non-epileptic hippocampi. A subsequent detailed computational simulation of a hippocampal circuit predicted that this aberrant theta-gamma coupling in DEE-13 is driven by increased persistent sodium currents (which is typical of *p*.*SCN8A* gain-of-function mutations [14–19]), and by selective loss of Oriens-lacunosum/moleculare (O-LM) cells and increased numbers of pyramidal cells. Immunohistochemistry and single-nucleus (sn)RNA sequencing confirmed that DEE-13 hippocampal assembloids selectively lose putative O-LM cells and have more excitatory neurons than CRISPR-corrected controls, with similar changes only in excitatory neurons in Cx+GE assembloids. Our findings highlight the importance of studying brain region-specific effects of genetic perturbations, and reveals interneuron-associated hippocampal network dysfunction in DEE-13. Our findings also elucidate possible distinct causal network pathologies in cortex and hippocampus underlying epilepsy versus cognitive impairments in DEE-13 and introduce hippocampal assembloids as a powerful platform for studying neural network development and pathology in the human hippocampus.

## 2 Results

### 2.1 Generation of Integrated Neural Assembloids

Cortical and hippocampal circuits in vivo contain both excitatory and inhibitory neural populations, with inhibitory neurons originating from the ganglionic eminences (GE). Organoids were directed toward GE fate using Sonic hedgehog (Shh) pathway agonists [20], a hippocampal fate using BMP4 and CHIR 99021 (a GSK3 inhibitor/Wnt agonist) [9], or a cortical fate through the absence of these patterning factors [20]. Without Shh, BMP4, or Wnt signaling, DEE-13 patient iPSC-derived organoids with a gain-of-function variant (p.R1872*>*L) [15] in the *SCN8A* gene (Mut) and matched CRISPR-corrected isogenic controls (iCtrl) showed cortical characteristics by day 56, expressing the cortical radial glial progenitor marker PAX6 [21], intermediate cortical progenitor marker TBR2 (EOMES) [22], and deep cortical layer marker CTIP2 (BCL11B) [23] (Fig. 1a). By day 120, these organoids expressed the superficial cortical layer marker SATB2 [24], and the deep cortical plate markers TBR1 [22] and CTIP2 (Fig. 1a). With Shh signaling but without BMP4 or Wnt signaling, both Mut and iCtrl organoids expressed markers typical of GE progenitors such as NKX2-1 [25] and OLIG2 [26], and the migratory interneuron marker DLX1 [27] by day 56, and by day 120 expressed the GABAergic inhibitory neuron marker GAD65 (GAD2) [28] (Fig. 1b). With BMP4 and Wnt signaling, and without Shh signaling, organoids at day 56 expressed the embryonic hippocampal marker NRP2 [29], as well as the granule cell markers PROX1 [30] and ZBTB20 [**?**] Fig. 1c). By day 120, these organoids expressed the cornu ammonis (CA)3 pyramidal cell marker KA1 (GRIK4) [31, 32] and the dentate and CA1/CA2 granule cell marker CTIP2 [33] (Fig. 1c and 1d).

**Figure 1.**
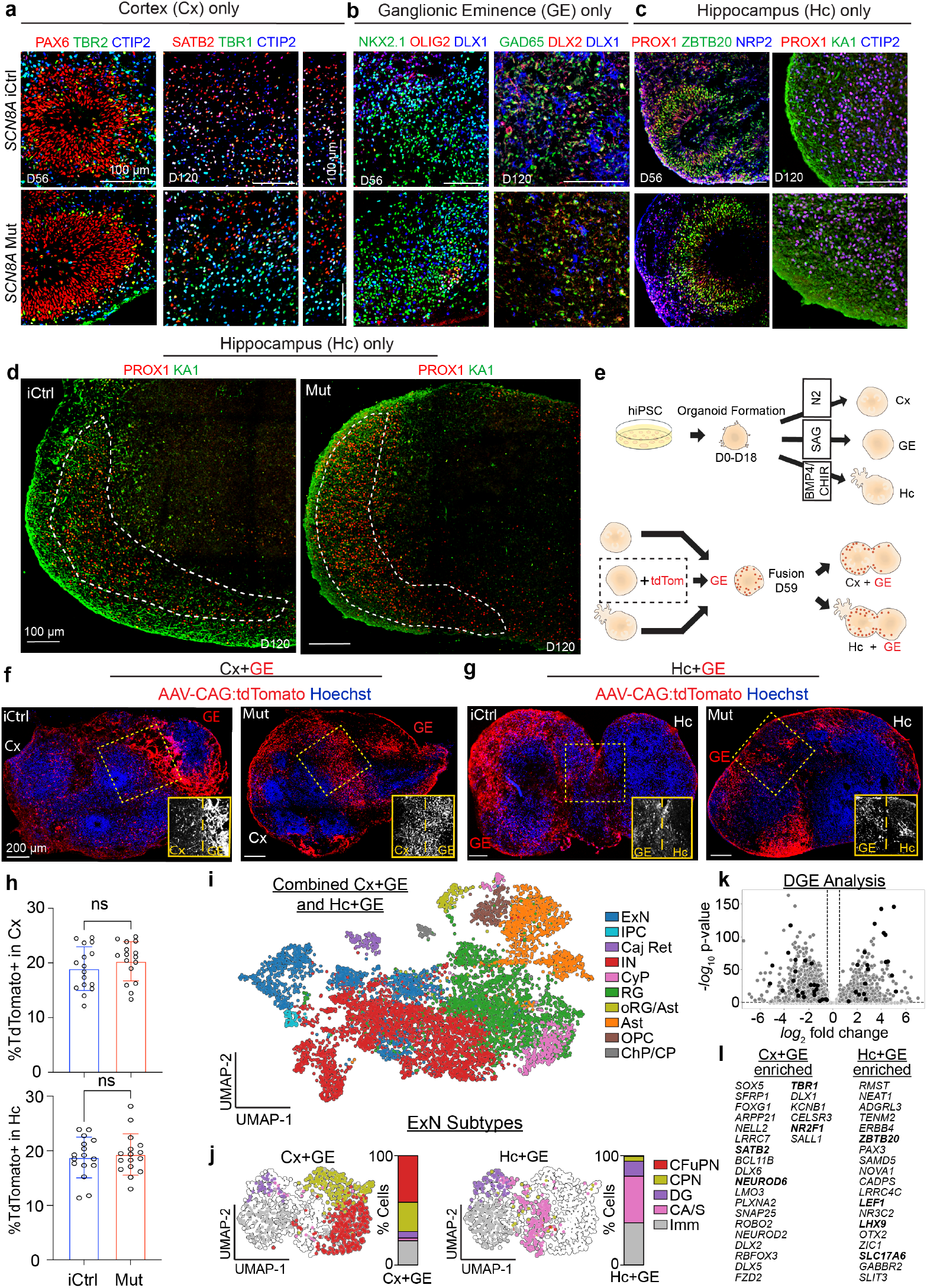
Generation and Characterization of Cortical and Hippocampal Assembloids: **a**, Immunohistochemical analysis of SCN8A patient iPSC-derived isogenic control (iCtrl) or mutant (Mut) unfused Cx organoids at specified ages. Both iCtrl and Mut organoids show similar formation of neural progenitors (PAX6, TBR2), deep and superficial neurons (CTIP2, TBR1, SATB2), and layered segregation of progenitors and neurons. **b**, iCtrl and Mut organoids display similar levels of GE progenitor and migratory interneuron markers (NKX2-1, OLIG2, DLX1, DLX2) and GABAergic interneuron marker GAD65 by D120. **c**, Immunohistochemical analysis of *p*.*SCN8A* iCtrl and Mut unfused hippocampal organoids at D56 and D120 shows expression of hippocampal neuron markers (ZBTBT20, NRP2) and dentate granule cells (PROX1), with CA3 and dentate granule layers at D120. **d**, Larger section showing PROX1-expressing dentate granule cells segregated from KA1-expressing CA3 region. **e**, Schematic of organoid generation, patterning, and fusion to generate assembloids. GE organoids treated with AAV1-CAG virus 3 days before fusion. **f**,**g**, At day 84 in vitro, iCtrl and Mut Cx+GE (f) and Hc+GE (g) show comparable migration of GE-derived Tdtomato-labeled cells. **h**, Quantification of Tdtomato+ cells shows no significant differences; n=4 independently generated assembloids per genotype, 4 sections per assembloid with ≥ 1,188 cells/sample for Cx+GE and 790 cells/sample for Hc+GE. p=0.6756 for Cx+GE and p=0.3254 for Hc+GE, student’s t-test. **i**, Integrated UMAP of day 120 iCtrl Cx+GE and Hc+GE with cell types. **j**, Subclustering excitatory neurons reveals unique subtypes with expected expression patterns. **k**,**l**, Volcano plot (k) and list (l) of differentially expressed genes (in order of statistical significance) showing unique genes distinguishing Cx+GE from Hc+GE (canonical genes in bold).

On Day 59, GE organoids were fused with either Cx or Hc organoids (Fig. 1e), and, to ensure integration of excitatory and inhibitory populations, a subset of GE organoids was treated with adeno-associated virus (AAV) CAG:tdTomato before fusion, showing widespread migration of tdTomato+ GE cells into adjacent Cx (Fig. 1f) and Hc compartments (Fig. 1g). At 2 weeks post-fusion, roughly 18% of cells in both Hc and Cx compartments were tdTomato positive, with no difference between Mut and iCtrls (Fig. 1h). By day 120, Cx+GE assembloids displayed intermixed neuronal populations, indexed by expression of the superficial cortical layer marker BRN2 (POU3F2) [34, 35] alongside GAD65+ interneurons, and the astrocyte marker GFAP [36] (Extended Data Fig. 1a). Similarly, by day 120, Hc+GE assembloids displayed diverse cell types including GFAP+ astrocytes and neurons (broadly marked by MYT1L expression). Much like unfused Hc organoids (Fig. 1d), the integrated Hc+GE assembloids maintained distinct dentate granule-like and CA3-like regions, marked by the spatial separation of PROX1 and KA1 expression (Extended Data Fig. 1b).

To further delineate compositional distinctions between cortical and hippocampal assembloids, we performed snRNA sequencing (snRNAseq) on iCtrl and Mut Cx+GE and Hc+GE assembloids at day 120. Both Cx+GE and Hc+GE assembloids demonstrated rich cell-type diversity and widespread *p*.*SCN8A* expression across conditions and cell types (Fig. 1i).

Cortical and hippocampal assembloids also expressed unique and distinguishing markers. Subtype analysis of excitatory neuronal markers revealed the predominance of corticofugal projection neurons (CFuPNs) and callosal projection neurons (CPNs) in Cx+GE versus markers of dentate gyrus (DG) and CA/subiculum (CA/S) in Hc+GE (Fig. 1j). In iCtrl Cx+GE assembloids at day 120, we observed enrichment of cortical markers (e.g. *SATB2, NEUROD6, TBR1*), and in Hc+GE iCtr, we saw enrichment of canonical hippocampal markers (e.g. *ZBTB20, LEF1, LHX9*) (Fig. 1k and 1l). The gene expression differences between Cx+GE and Hc+GE assembloids were consistent with what has been previously reported in the developing mammalian cortex and hippocampus, respectively [37, 38].

### 2.2 DEE-13 Cortical Assembloids are Hyperexcitable

To evaluate circuit-specific effects of gain-of-function *p*.*SCN8A* mutations in DEE-13, we performed extracellular recordings of local field potentials (LFPs) and two-photon microscopy-based calcium imaging of Cx+GE and Hc+GE assembloids. We used validated iPSCs from two DEE-13 patients with unique *p*.*SCN8A* variants [15]. Patient 1 (p.R1872*>*L) had seizure onset at 1 month with intellectual disability and medically refractory seizures. Patient 2 (p.V1592*>*L) had seizure onset at 5 months with developmental delay and well-controlled epilepsy on medication [15]. These were compared to a CRISPR-corrected isogenic control for Patient 1. Replicating our prior results [4], iCtrl Cx+GE assembloids generated oscillatory LFPs at frequencies similar to those observed in the cortex in vivo (Fig. 2a). Both Mut Patient 1 (P1) and Patient 2 (P2) Cx+GE generated LFPs with intermixed bursts of high-amplitude discharges and periods of quiescence (Fig. 2b-c). Mut P1, but not Mut P2, Cx+GE assembloids had significantly more sustained high-amplitude LFP activity compared to iCtrls (arrow and bracket in 2b and Fig. 2d), consistent with P1’s more severe epilepsy [15]. Quantification of LFP discharges into clinically defined spikes, sharp-waves, and sustained “long duration discharges” (LDDs) revealed a significant increase in spikes and LDDs in Mut P1 Cx+GE, along with a non-significant but substantial increase in sharp waves. It also revealed a significant increase in spikes in Mut P2 Cx+GE assembloids (Fig. 2e), without concomitant increases in sharp waves or LDDs, consistent with P2’s less severe clinical phenotype.

**Figure 2.**
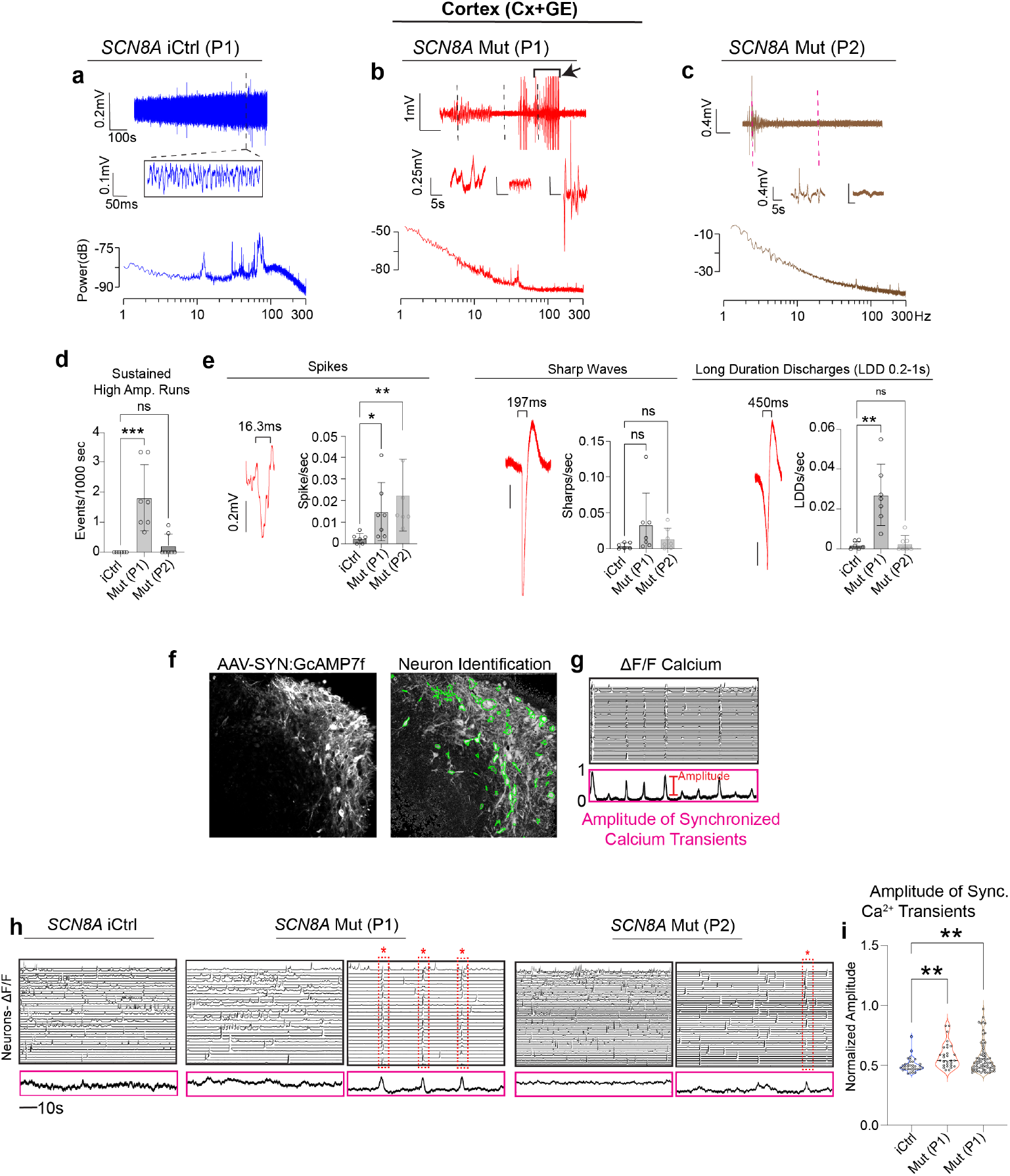
DEE-13 Cortical Assembloids Demonstrate Network Hyperexcitability: **a**, Raw trace (top) and time expansion (middle) from an LFP recording of an isogenic control (iCtrl, P1) *p*.*SCN8A* cortex assembloid (Cx+GE). Bottom, periodogram from the entire raw trace. **b**, Raw trace (top) and three time expansions (middle) from the LFP recording of a Mut P1 (p.R1872*>*L) *p*.*SCN8A* Cx+GE assembloid, with time expansions near the dotted lines in the raw trace. Bottom, periodogram from the entire raw trace. **c**, Raw trace (top) and two time expansions (middle) from the LFP recording of a Mut P2 (p.V1592*>*L) *p*.*SCN8A* Cx+GE assembloid, with time expansions near the dotted lines in the raw trace. Bottom, periodogram from the entire raw trace. **d**, Number of sustained high amplitude events/1000s in iCtrl and Mut, defined as sustained runs ≥30s with RMS amplitude ≥ 5x greater than the preceding 30s. **e**, Frequency of spikes, sharp waves, and LDDs from iCtrl and Mut Cx+GE LFP recordings, with a representative example of each shown in red. For **d** and **e**, n=6 biologically independent samples for iCtrl, n=7 for Mut P1 and P2. Kruskal-Wallis test with Dunn’s multiple comparisons test. ***p=0.0006, *p=0.0179, **p=0.0030 (spikes), **p=0.0079 (LDDs). **f**, Schematic for calcium transients identification and functional clusters generation after multiphoton imaging with a genetically encoded calcium indicator. **g**, Normalized *δ*F/F of calcium indicator activity in iCtrl Cx+GE. Each line represents a single neuron’s activity. **h**, Normalized *δ*F/F of calcium indicator activity in iCtrl, Mut P1, and Mut P2 Cx+GE. Decorrelated traces with minimal synchrony (left) and highly synchronous discharges (right) in Mut Cx+GE. **i**, Significant increase in average amplitude of Ca2+ transients in Mut P1 and P2 Cx+GE vs. iCtrl. Linear mixed effects model. **p=0.0041 (iCtrl vs Mut P1), **p=0.0088 (iCtrl vs Mut P2).

To measure single-cell activity, we infected Cx+GE assembloids with AAV1 Syn:GCaMP7f virus (Fig. 2f) and recorded spontaneous calcium activity as changes in GCaMP7f fluorescence (ΔF/F) two weeks later. To quantify network-wide hypersynchronization, associated with an “epileptic-like” hyperexcitable state, we calculated the percentage of cells in each frame that were simultaneously active (Fig. 2g). Network-wide activity was largely desynchronized in iCtrl Cx+GE assembloids (Fig. 2h), whereas both Mut P1 and Mut P2 Cx+GE assembloids displayed periods with asynchronous activity interspersed with periods showing network-wide synchronization (Fig. 2h, Supplementary Videos 1-5). The amplitude of synchronized calcium transients, quantified as the percentage of active cells per frame, was significantly increased in both Mut P1 and Mut P2 Cx+GE assembloids relative to iCtrl (Fig. 2i).

Unlike Cx+GE assembloids with gain-of-function *p*.*SCN8A* mutations, LFP recordings from Hc+GE Mut assembloids did not display overt hyperexcitability. No high-amplitude bursts or spikes were observed in LFPs recorded from Hc+GE assembloids derived from either DEE-13 patient’s iPSCs, similar to iCtrl Hc+GE assembloids (Fig. 3a). Power spectra for iCtrl and Mut P1 showed sustained oscillations lasting at least five minutes (Fig. 3b) and spectral power quantification revealed no consistent differences in low-frequency bands (Extended Data Fig. 2a). Mut P2 LFPs also showed oscillations at multiple frequencies, but these lasted about one minute at a time (inset, Fig. 3b).

**Figure 3.**
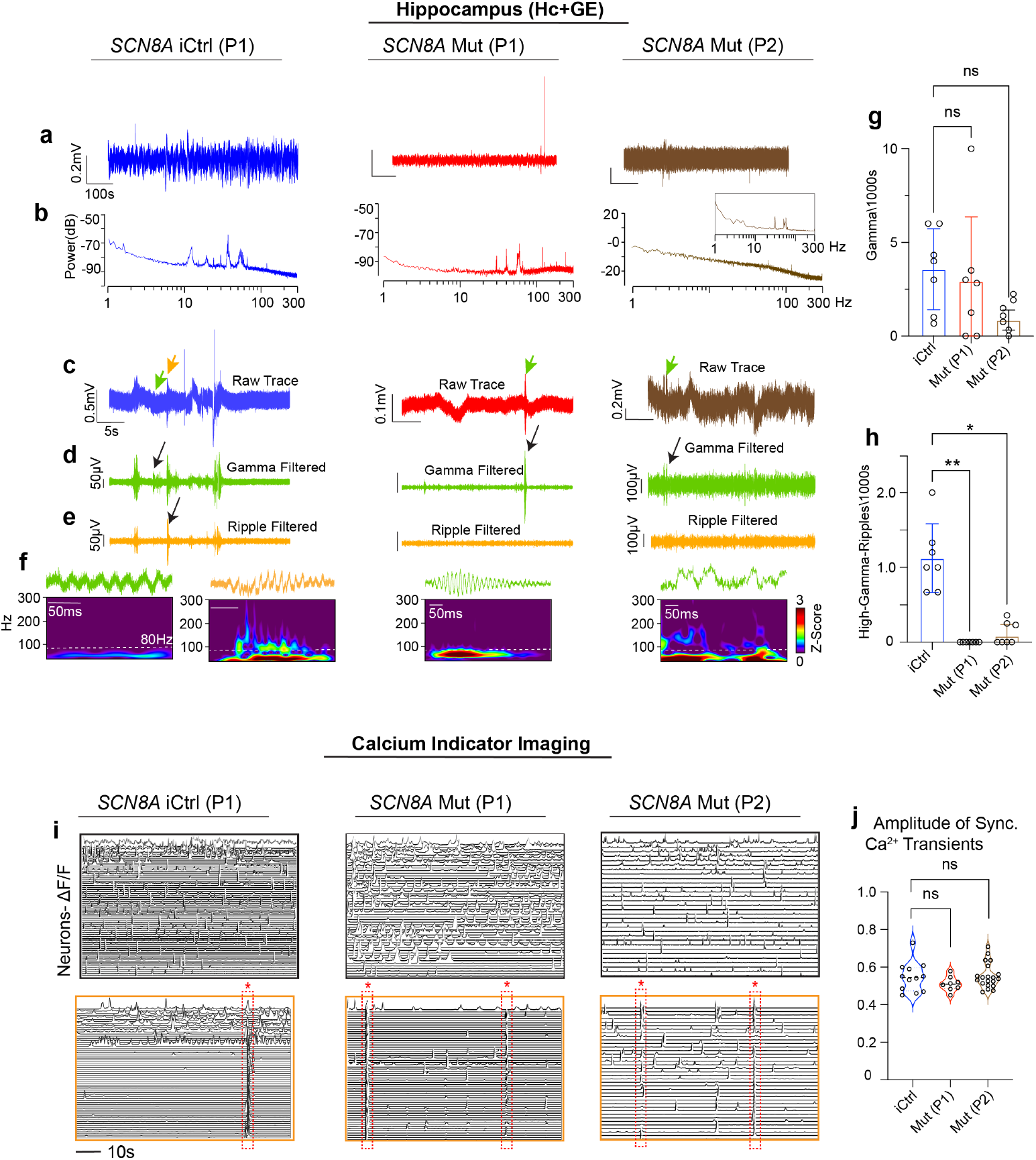
DEE-13 Hippocampal Assembloids Show a Reduction in Fast Oscillations but Lack Overt Hyperexcitability: **a**, Raw traces from LFP recordings of iCtrl (P1, blue), Mut P1 (p.R1872*>*L, red), and Mut P2 (p.V1592*>*L, brown) *p*.*SCN8A* hippocampal fusion (Hc+GE) assembloids. **b**, Corresponding periodograms of the entire recording segment in (a). Mut P2 lacked sustained oscillatory activity ≥ 300s, as seen in iCtrl and Mut P1, but showed shorter bouts (∼60s) of oscillations (inset periodogram). **c**, Time-expanded segments of traces in (**a**), with green arrows indicating putative gamma bursts and an orange arrow indicating a high-gamma/ripple burst. **d**, Gamma oscillations (30-80 Hz) from segments in (**c**), color-coded in green, with one gamma run (black arrow). **e**, High gamma/ripple frequency activity (80-160 Hz) from segments in (**c**), color-coded in orange, with a ripple (only in iCtrl, black arrow). Mut P1 and P2 lack peaks in ripple-filtered traces but show slower gamma activity. **f**, Morlet plots and time expansions of individual gamma and ripple discharges shown by color-coded arrows in (**c**), and black arrows in (**d**) and (**e**). **g**, Quantification of gamma episodes/1000s reveals no differences between groups. **h**, Quantification of high-gamma/ripples/1000s shows significantly more high-frequency activity in iCtrl vs. Mut P1 and P2. n=7 independently generated assembloids/group. 1-way ANOVA with Dunnett’s multiple comparison test, **p=0.0042, *p=0.0222. **i**, Representative ΔF/F plots from calcium indicator imaging of iCtrl P1, Mut P1, and Mut P2 Hc+GE assembloids. iCtrl and Mut assembloids show decorrelated (top) and synchronized (bottom, dotted red boxes) activity. Quantification of synchronization amplitude shows no significant differences between groups. n=6 independently generated assembloids/group. A linear mixed effects model was used.

Given the absence of clear hyperexcitability, we evaluated if iCtrl Hc+GE assembloids generated canonical hippocampal circuit activities and if these were altered in *p*.*SCN8A* variant-expressing assembloids. Sharp wave ripples (SWRs), high-frequency oscillations riding on a high-amplitude discharge, are well-established hippocampal activities associated with memory consolidation [39, 40]. We identified all discharges in raw LFP traces and filtered traces in the low gamma (30-80 Hz) (Fig. 3d) and high gamma/ripple (80-160 Hz) (Fig. 3e) ranges. Despite observing gamma epochs in all Hc+GE lines (Fig. 3d), including gamma frequency discharges (green and black arrows in Fig. 3c and 3d), high gamma/ripple epochs were almost exclusively in iCtrl Hc+GE (Fig. 3e) and absent from the Mut specimens. Morlet plots of individual discharges confirmed solely low gamma frequency discharges in Mut P1 and P2, and additional higher frequency ripples in iCtrl, though none of these resembled classic SWRs (Fig. 3f). Quantification showed no significant changes in low gamma activity between lines (Fig. 3g), but did show notable reductions in high-gamma-ripple activity in Mut P1 and P2 Hc+GE assembloids compared to iCtrl (Fig. 3h).

Similar to initial LFP results (Fig. 3a-b), calcium indicator data were similar between iCtrl, Mut P1, and Mut P2, with all lines showing a mix of synchronous and asynchronous activity (Fig. 3i, Supplementary Videos 6-11). Unlike for cortex, quantification revealed no significant differences in synchronized events beyond the intrinsic mixture seen in iCtrl Hc+GE assembloids (Fig. 3j).

### 2.3 Aberrant Network Activity in DEE-13 Hc+GE Assembloids Resembles Disruptions Seen in Epilepsy Patients

To further explore hippocampus-associated network dynamics in iCtrl Hc+GE and their perturbations in Mut specimens, we examined coupling between hippocampal theta (3-10 Hz) phase and gamma/high gamma (30-120 Hz) amplitude oscillations (Fig. 4a). Theta-gamma phase-amplitude coupling (PAC) is crucial in the mammalian hippocampus for encoding, maintenance, and retrieval of information [41]. While there was no difference in theta-gamma coupling strength between iCtrl and Mut Hc+GE assembloids (Extended Data Fig. 2b), the pattern of coupling was “monophasic” in iCtrl, i.e., the entire range of gamma amplitudes locked to the same theta phase, typically the peak (Fig. 4b). Conversely, theta-gamma PAC was disordered in both P1 and P2 Mut Hc+GE, with different gamma amplitudes coupling to various theta phases (Fig. 4c-d). The monophasicity of theta-gamma coupling was significantly reduced in both P1 and P2 Mut Hc+GE assembloids (Fig. 4e), despite no difference in theta power (Fig. 4f) or gamma power (Extended Data Fig. 2a). We evaluated whether theta oscillations in Mut Hc+GE assembloids were unstable rather than reduced in power, potentially compromising continuous monophasic phase-locking. Using the K-statistic of the 0-1 chaos test [42, 43], we found that theta oscillations were significantly less stable in both Mut Hc+GE assembloids relative to iCtrls (Fig. 4g). Corroborating this, we observed more unstable single-cell spiking patterns in both Mut samples relative to iCtrl Hc+GE assembloids (Fig. 4h).

**Figure 4.**
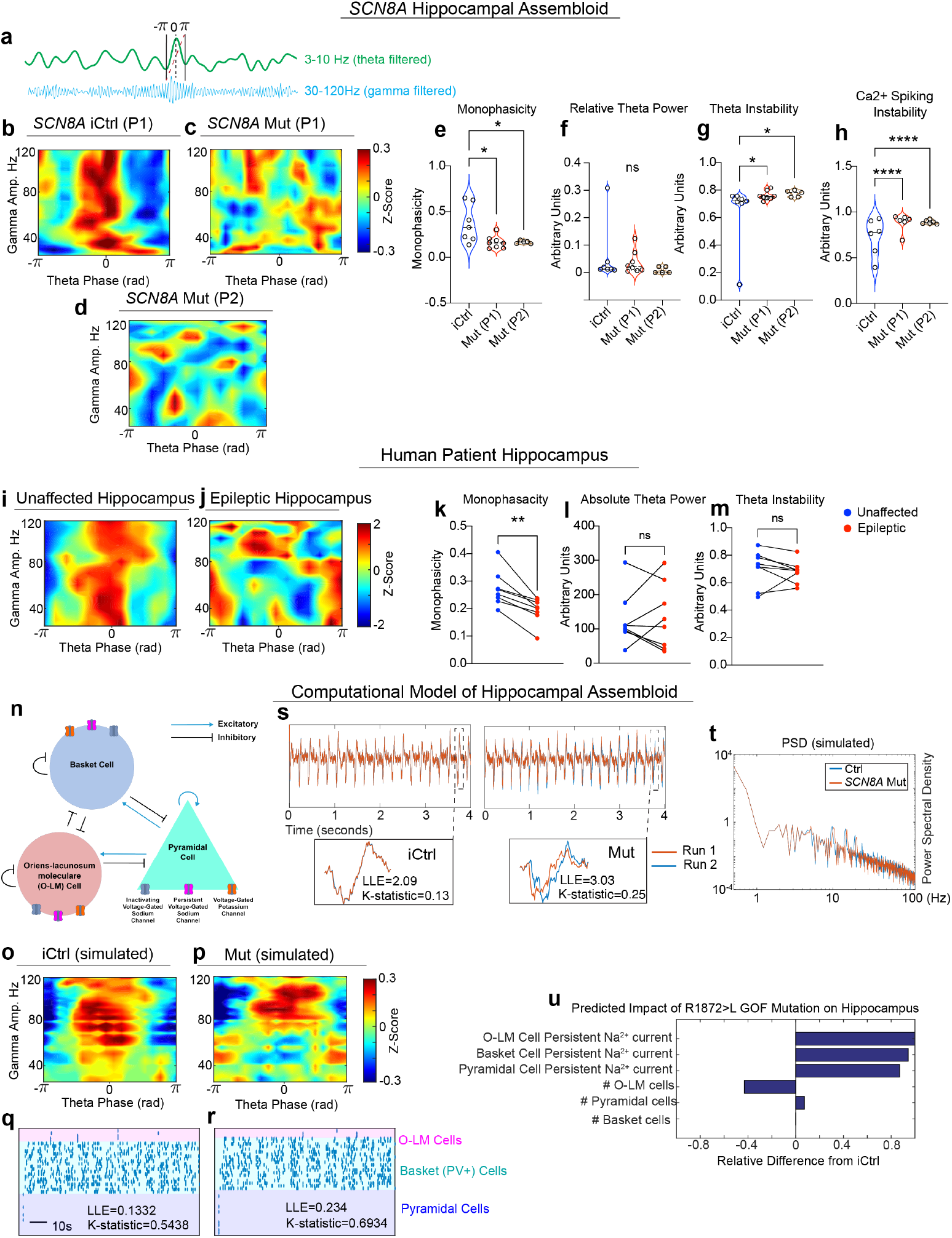
Perturbation of Theta-Gamma Coupling in Both DEE-13 Hippocampal Assembloids and Human Temporal Lobe Epilepsy: **a**, An example of phase-amplitude coupling (PAC) in an iCtrl Hc+GE assembloid showing higher gamma amplitude coupled to the 0 phase of theta (3-10 Hz). **b-d**, Heatmaps of theta phase (x-axis) and gamma amplitude (y-axis) coupling in *p*.*SCN8A* iCtrl (b), Mut (c, d) Hc+GE assembloids. iCtrl shows monophasic PAC, Mut shows disordered PAC. **e-h**, Quantification of monophasicity (e), theta power (f), theta instability (g), and Ca^2+^ spiking instability (h) in iCtrl and Mut Hc+GE assembloids. Monophasicity reduced in Mut (one-tailed Wilcoxon rank-sum test), no significant differences in theta power (two-tailed Wilcoxon rank-sum test, n=5-8 independently generated assembloids/genotype). Mut shows increased theta and Ca^2+^ spiking instability (linear mixed effects model, n=6 assembloids/line). **i-j**, PAC heatmaps in the healthy (i) and epileptic (j) hippocampus of one patient. Healthy shows monophasic coupling, epileptic shows disordered coupling. **k-m**, Monophasicity (k), theta power (l), theta instability (m) in healthy and epileptic hippocampi of 8 temporal lobe epilepsy patients. Monophasicity reduced in epileptic hippocampi (one-tailed Wilcoxon ranksum test), no differences in theta power/instability (two-tailed Wilcoxon rank-sum test). **n**, In silico CA3 model simulating iCtrl and Mut Hc+GE electrophysiology. **o-r**, Simulated gamma amplitude vs. theta phase heatmaps (o-p), raster plots of cell spiking (q-r) for iCtrl and Mut Hc+GE assembloids. **s**, LFPs from iCtrl and Mut simulations showing greater divergence in Mut. **t**, Power spectral density plots for simulated iCtrl and Mut Hc+GE assembloids showing no marked changes in theta or gamma power. **u**, Predicted impact of R1872*>*L *p*.*SCN8A* mutation on hippocampus.

To assess the in vivo relevance of these observed PAC deficits, we analyzed theta-gamma coupling in bilateral intracranially recorded LFPs from the hippocampi of 6 eight temporal lobe epilepsy (TLE) patients during task-free and seizure-free periods. Remarkably, we found reduced monophasicity in the theta-gamma coupling of each patient’s epileptic hippocampus relative to their healthy hippocampus (Fig. 4i-k). Together, these results suggest that loss of monophasic theta-gamma coupling, first observed in our Hc+GE assembloids, may be a hallmark of altered hippocampal circuit function in epilepsy. Like our Mut Hc+GE assembloids, there was no change in theta power in the patients’ epileptic hippocampi (Fig. 4l). However, unlike our Mut assembloids, there was no clear increase in theta oscillation instability (Fig. 4m), potentially suggesting different mechanisms for loss of theta-gamma monophasicity in TLE compared to DEE-13.

### 2.4 In Silico Modeling Predicts Changes in Cell Numbers

To predict mechanisms underlying the electrophysiological phenotype of Mut Hc+GE assembloids, we developed an in silico CA3 circuit simulation extending a previous model [44]. This model included fast-spiking basket cells, slow-spiking O-LM cells, and pyramidal cells, each described by Hodgkin-Huxley equations (see Methods) (Fig. 4n). We introduced persistent sodium currents into the circuit equation for each neuron type, reflecting effects of the specific gain-of-function *p*.*SCN8A* variants on non-inactivating sodium conductance [14–19].

For our simulation of iCtrl Hc+GE assembloids, we set cell proportions and numbers based on preliminary immunohistochemistry data. Using genetic optimization, we modified all other model parameters to generate monophasic theta-gamma coupling, as observed in Hc+GE assembloids and healthy human hippocampus (Fig. 4o). We then evaluated cell type-specific changes driving electrophysiological phenotypes in *p*.*SCN8A* Mut Hc+GE assembloids. We used another genetic optimization algorithm to alter persistent sodium currents and the number of pyramidal, basket, and O-LM cells in the model, such that the model would recapitulate the Mut Hc+GE phenotype (namely, reduced monophasic theta-gamma coupling, destabilized single-cell spiking and circuit-level oscillatory dynamics, and minimal changes in theta or gamma power, Fig. 4o-t). The resulting Mut model showed large increases in persistent sodium currents for all cell types, a loss of O-LM cells, and increased pyramidal cells, with no changes in basket cells (Fig. 4u). Interestingly, dysfunction of somatostatin-expressing (SST) hippocampal interneurons, which include O-LM cells, has been demonstrated to disrupt hippocampal circuitry and impair memory in another neurodevelopmental disorder, Rett syndrome [45]. Moreover, when the persistent sodium current was renormalized in our in silico model, the monophasic theta-gamma coupling was only partially restored (Extended Data Fig. 3a), indicating that normal (iCtrl) cellular composition was necessary to maintain monophasic theta-gamma coupling in DEE-13 Hc+GE assembloids. Pathological changes in persistent sodium currents are consistent with known effects of gain-of-function *p*.*SCN8A* mutations [14–19]. While O-LM cell loss is observed in multiple temporal lobe epilepsy models [46–48], the cellular composition of hippocampal circuits in DEE-13 has not been evaluated to our knowledge.

### 2.5 *p*.*SCN8A* Hc+GE Assembloids Display Alterations in Interneuron and Excitatory Neuron Development

To asses whether the aberrant electrophysiology in *p*.*SCN8A* Mut Hc+GE assembloids is driven by changes in the balance of O-LM and pyramidal cells as predicted by our computational model, we used a combination of immunohistochemical (IHC) and snRNAseq approaches to evaluate the cellular composition of *p*.*SCN8A* iCtrl vs. Mut Cx and Hc assembloids. Based on the in silico prediction, we focused broadly on excitatory and inhibitory balance, and sought to identify the in vitro equivalent of in silico “basket” and O-LM interneurons in addition to excitatory neurons. We therefore quantified the expression of GAD65 together with Parvalbumin (PV), associated with fast-spiking/basket interneurons, and SST, since O-LM are known to express SST, as well as CAMKII-*α* (CAMK2A) to distinguish excitatory neurons. Mut P1 Hc+GE showed a significant reduction of SST+ but not PV+ interneurons, and an increase in CAMKII-*α*+ excitatory neurons compared to iCtrl (Fig. 5a and 5b), matching our computational model’s predictions of the key contributors to the in vitro loss of monophasic theta-gamma coupling in DEE-13 hippocampus. Mut P2 Hc+GE showed similar changes, with an additional reduction in PV+ interneurons (unlike Mut P1, Fig. 5a and 5b). The latter is consistent with some of the alternative in silico Mut simulations identified by a systematic parameter sweep (Extended Data Fig. 3b and 3c). Comparable IHC on Cx+GE assembloids showed no reduction in total (GAD65+) interneurons (Fig. 5c), and no significant changes in PV (Fig. 5c). However, there was a significant increase in CAMKII-*α*+ excitatory neurons in both P1 and P2 Mut Cx+GE compared to iCtrl Cx+GE (Fig. 5d).

**Figure 5.**
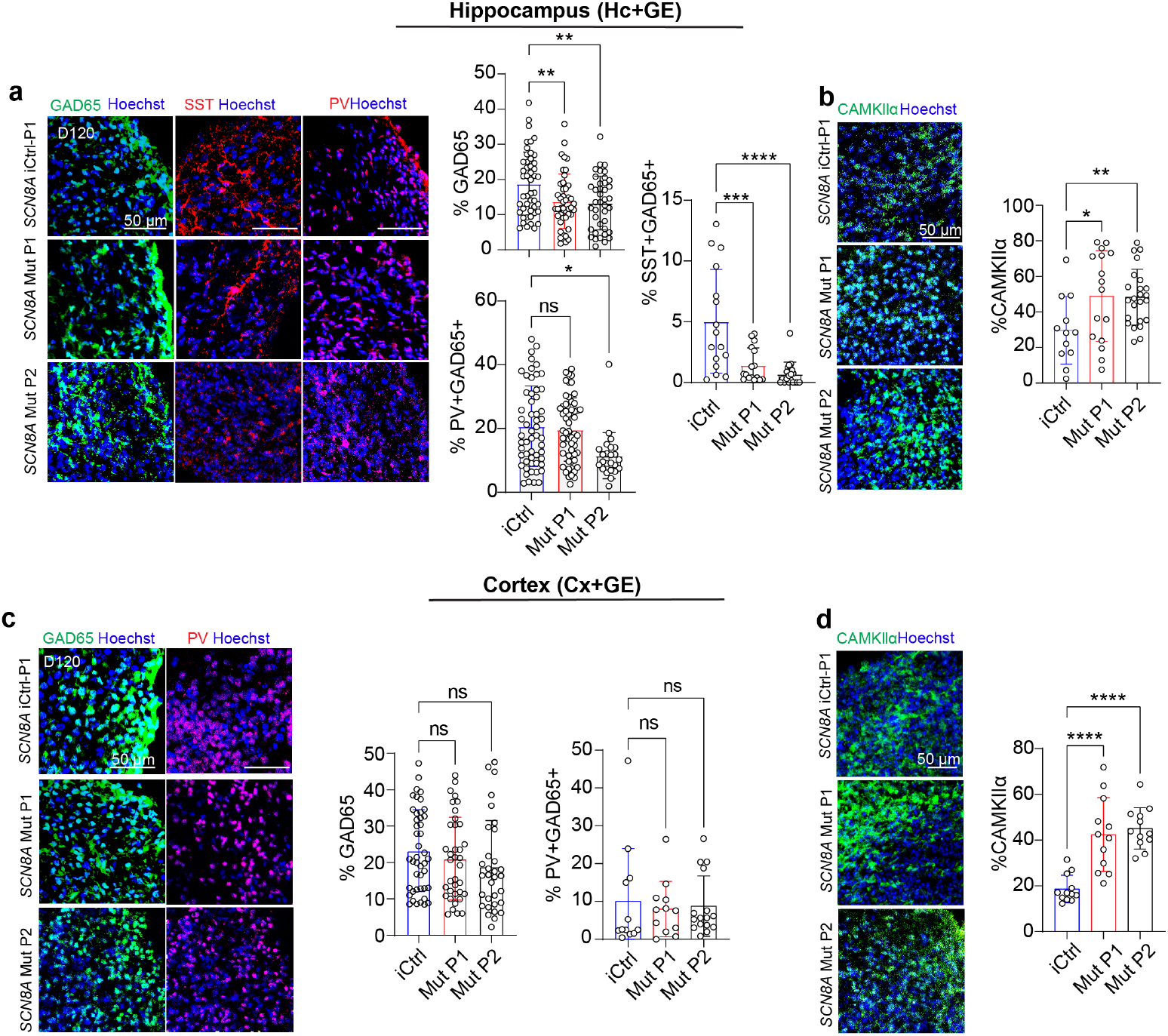
Hippocampal but not Cortical DEE-13 Assembloids Demonstrate Reduced Inhibitory Interneuron Numbers: **a**, Immunohistochemical (IHC) analysis of Hc+GE assembloids reveals a reduction in the pan-inhibitory interneuron marker glutamic acid decarboxylase-65 (GAD65/GAD-2) and in the somatostatin (SST) subtype of interneurons in both Mut P1 and Mut P2 compared to iCtrl. IHC revealed a significant reduction in parvalbumin+ (PV+) interneurons in Mut P2, but no significant change in Mut P1. **b**, IHC analysis for excitatory neurons with CAMKII-*α* revealed a significant increase in both Mut P1 and Mut P2 compared to iCtrl. n=3-6 independently generated assembloids per genotype, 4 sections per assembloid with ≥4,280 cells per sample. For Mut P1 and Mut P2 respectively; GAD65 **P=0.0037 and **p=0.0019; PV *p=0.0136; SST ***p=0.00016 and ****p=4.24E-6; CAMKII-*α* *p=0.0188 and **p=0.0088. **c**, IHC for both GAD65 and PV in Cx+GE assembloids reveal no significant differences between iCtrl and Mut P1 or Mut P2. **d**, IHC for CAMKII-*α* reveals a significant increase in both Mut P1 and Mut P2 compared to iCtrl. n=3-11 independently generated assembloids per genotype, 4 sections per assembloid with ≥ 2681 cells per sample. For Mut P1 and Mut P2 respectively; CAMKII-*α* ****p=7.53E-7 and ****p=1E-7.

### 2.6 Transcriptomic Analysis Reveals Selective Hc+GE Interneuronopathy in *p*.*SCN8A* Mut Hc+GE Assembloids

To further explore the divergent developmental trajectories between Cx+GE and Hc+GE R1872*>*L *p*.*SCN8A* assembloids and identify neuronal subtype differences, we used an expanded snRNAseq dataset that included both iCtrl and Mut P1 Cx+GE and Hc+GE assembloids at day 120, as well iCtrl and Mut P1 Hc+GE at day 84. This analysis confirmed and expanded on our IHC findings and computational predictions. Cx+GE assembloids consisted of diverse but expected cell types (Fig. 6a and Extended Data Fig. 4-6). Consistent with IHC results, Mut Cx+GE assembloids showed a relative increase in excitatory neurons (ExN, Fig. 6b) without significant changes in other major cell populations, including interneurons (INs, Fig. 6b). No substantial changes were seen in excitatory neuron subtypes between iCtrl and Mut (Fig. 6c). Subtype analysis of Cx+GE inhibitory interneurons showed an increase in mixed CGE/LGE-like cells in Mut assembloids (Fig. 6d).

**Figure 6.**
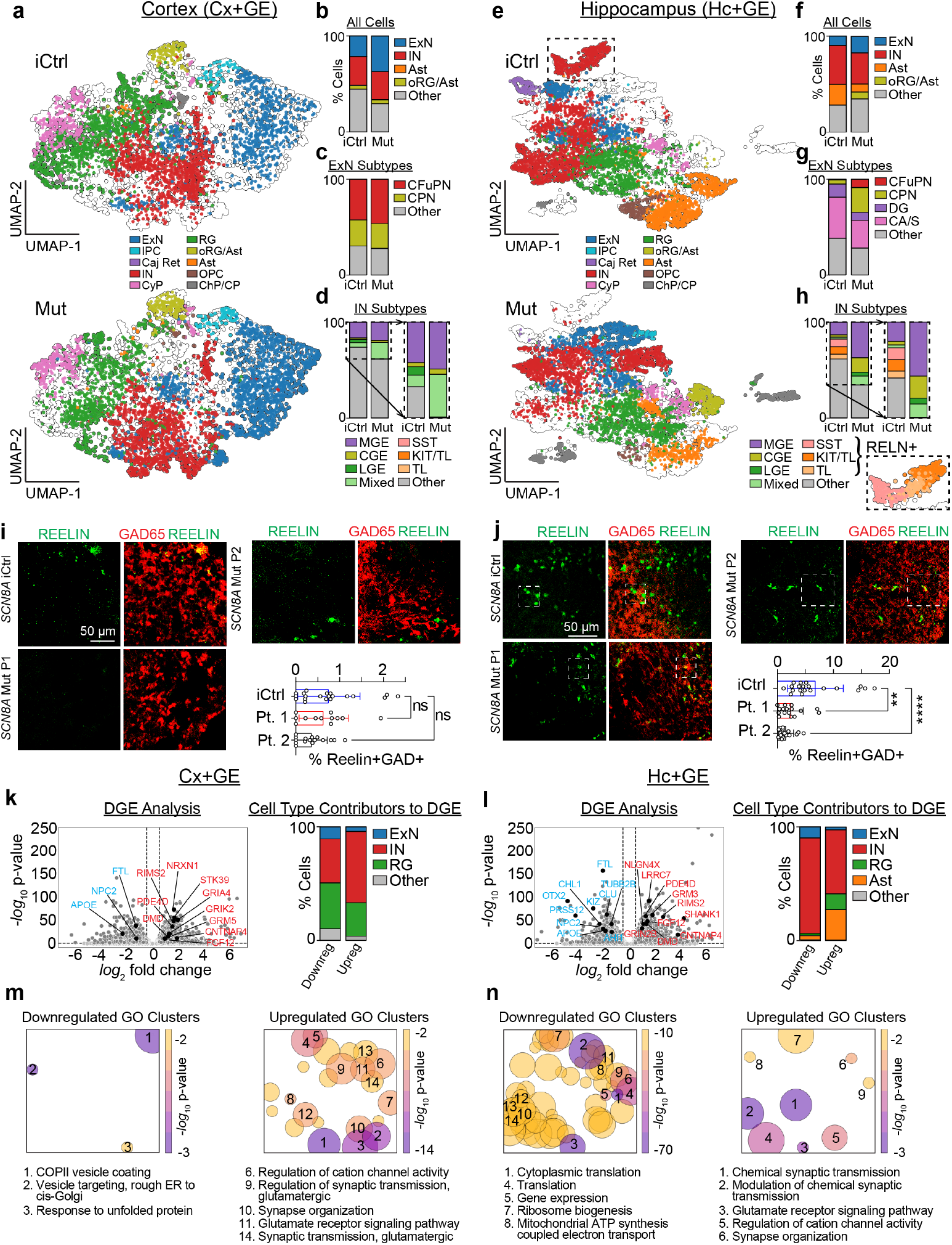
DEE-13 Differentially Affects Circuit Composition and Gene Expression in Cortical Versus Hippocampal Assembloids: **a**, 2-D UMAP of cells from iCtrl and Mut P1 Cx+GE assembloids. **b**, Mut P1 Cx+GE assembloids show more excitatory neurons (ExN) compared to iCtrl. **c**, No significant changes in ExN subtypes (CFuPN, CPN) between Mut P1 and iCtrl Cx+GE. **d**, Mut P1 Cx+GE assembloids show more mixed CGE- and LGE-like inhibitory interneurons (INs) compared to iCtrl. **e**, UMAP of cells from iCtrl and Mut P1 Hc+GE assembloids. **f**, Mut P1 Hc+GE assembloids show more ExNs, fewer INs, fewer astrocytes (Ast), and more oRG/Ast compared to iCtrl. **g**, Mut P1 Hc+GE assembloids show more cortex-like CFuPN and CPN ExNs and fewer hippocampal DG-like and CA/S-like ExNs compared to iCtrl. **h**, Mut P1 Hc+GE assembloids lack *RELN* + INs present in iCtrl Hc+GE. **i**,**j**, IHC shows few cells coexpressing Reelin and GAD65 in cortical assembloids across iCtrl, Mut P1, and Mut P2 lines, while hippocampal assembloids show a substantial population in iCtrl that is lost in Mut lines (*p* < .0001, linear mixed effects model, 5 iCtrl, 4 Mut P1 and P2 assembloids, 4 sections per assembloid, ≥ 6,324 cells/sample). **k**,**l**, Differential gene analysis volcano plots of Cx+GE and Hc+GE, respectively, shows selected upregulated genes (red) in Mut involved in glutamatergic pathways and selected downregulated genes (blue) that are associated with epilepsy and intellectual disability when their expression is reduced and are shown in relative proportion. **m**,**n**, Gene ontology (GO) analysis of Cx+GE and Hc+GE, respectively, shows upregulation of glutamatergic pathways and downregulation of basic cellular tasks in hippocampal Mut P1 assembloids. Larger circles correspond to more GO terms. **Additional Abbr**. IPC = intermediate progenitor cells, Caj Ret = Caja-Retzius cells, CyP = cycling progenitors, RG = radial glia, OPC = oligodendrocyte precursor cells, ChP/CP = choroid plexus/cortical plate, MGE = medial ganglionic eminence-like INs.

The hippocampal assembloids also consisted of diverse cell types (Fig. 6e and Extended Data Fig. 4-6), with more substantial differences between Mut and iCtrl samples. These changes were consistent with and expanded on our IHC findings. Mut Hc+GE showed a relative increase in excitatory neurons (ExN, Fig. 6f) consistent with IHC and in silico modeling predictions. Unlike any other assembloid including iCtrl Cx+GE, iCtrl Hc+GE exhibited a well-defined astrocyte population (i.e. *GFAP* +,*AQP4* +, *S100B* +). In Mut Hc+GE, the proportion of these astrocytes decreased concomitant with an increase in the mixed outer radial glia/astrocyte co-cluster that was prominent in all Cx+GE assembloids (Ast, oRG/Ast, Fig. 6f). Cajal-Retzius cells were also largely exclusive to iCtrl Hc+GE. Subtype analysis of excitatory clusters revealed that changes in Mut Hc+GE were driven by increased proportions of excitatory cells expressing cortical-like deep and upper layer markers (CFuPN and CPN), with a relative reduction in excitatory neurons expressing hippocampal DG-like and CA/S-like markers (Fig. 6g). These findings suggest that the gain-of-function *p*.*SCN8A* mutation may drive hippocampal cell identities toward a cortical-like fate.

The iCtrl Hc+GE assembloids also had a unique *RELN*-expressing interneuron cluster (boxed cluster, Fig. 6e) that was absent in Mut Hc+GE (and all Cx+GE). As a result, and unlike Cx+GE, Mut Hc+GE had an overall reduction in inhibitory interneurons (INs, Fig. 6f). Subtype analysis showed that the *RELN*-expressing interneuron cluster selectively reduced in Mut Hc+GE included *SST* (putative OLM) interneurons and *CHRM2*-expressing trilaminar (TL) interneurons (Fig. 6h). The reduction in putative O-LM/SST interneurons is consistent with our in silico prediction (Fig. 4) and IHC results (Fig. 5). The reduction in TL interneurons is also consistent with our electrophysiologic observations, as TL interneurons are highly associated with ripple activity [49], and we recorded a significant reduction in high gamma/ripple oscillations in Mut Hc+GE (Fig. 3). Subsequent IHC confirmed limited numbers of Reelin+ interneurons in Cx+GE (Fig. 6i) and showed that this population was present in iCtrl Hc+GE but significantly reduced in both P1 and P2 Mut Hc+GE. Notably, snRNAseq analysis from both iCtrl and Mut day 84 Hc+GE demonstrated the absence of this *RELN*-expressing cluster (Extended Data Fig. 7), despite other hippocampal markers being present by this time. This suggests that these are relatively late-developing IN populations in iCtrl Hc+GE that fail to develop in Mut Hc+GE.

Next we used differential gene expression (DGE) and gene ontology analysis to identify potential pathways underlying the striking cellular differences that result from *p*.*SCN8A* variant expression in hippocampus-vs. cortex-like assembloids. DGE analysis of upregulated genes showed increases in glutamatergic genes in both Cx+GE (Fig. 6k) and Hc+GE (Fig. 6l), suggesting a potential broad mechanism for hyperexcitability in DEE-13. Unexpectedly, the major cell type contributor to these upregulated genes was inhibitory interneurons with only a minor contribution from excitatory neurons (Fig. 6k and 6l). This suggests some degree of interneuron dysfunction and a possible shift away from excitatory-inhibitory balance. This upregulation in glutamatergic genes and shift toward a more excitatory state in both Mut Cx+GE and Hc+GE assembloids was supported by gene ontology analysis and super-clustering, showing over-representation of excitatory synapse and glutamate signaling pathways (Fig. 6m and 6n). In contrast, DGE analysis of downregulated genes in Mut Cx+GE and Hc+GE demonstrated regional-specific changes. For example, Mut vs. iCtrl Hc+GE demonstrated a larger depletion of genes important for normal development compared to Mut vs. iCtrl Cx+GE, and many of these genes are associated with epilepsy or intellectual disability [50] when dysfunctional (Fig. 6k and 6l). While in Mut Cx+GE the contribution to these downregulated genes was shared between multiple cell types including interneurons, radial glia, and excitatory neurons (Fig. 6k), in Mut Hc+GE the contribution was overwhelmingly from inhibitory interneurons (Fig. 6l). Gene ontology analysis further supported these findings, revealing few down-regulated pathways in Mut vs. iCtrl Cx+GE assembloids (Fig. 6m) and, conversely, many downregulated pathways involved in important cellular viability/housekeeping tasks (e.g., translation, gene expression, metabolism) in Mut vs. iCtrl Hc+GE assembloids (Fig. 6n). Together, these findings indicate that *p*.*SCN8A* results in two distinct changes where interneurons are the primary cellular contributor to pathology: 1) nonspecific upregulation of hyperexcitability-related genes, which may contribute to excitatory-inhibitory imbalance, and 2) a hippocampal-specific downregulation of certain genes involved in important basic cell functions that lead to phenotypic abnormalities when these genes are mutated.

## 3 Discussion

Our study used human brain assembloids to explore region-specific network effects of *p*.*SCN8A* gain-of-function variants in DEE-13. By generating cortical (Cx+GE) and hippocampal (Hc+GE) assembloids, we elucidated distinct neural dynamics and cellular disruptions caused by these variants in cortical versus hippocampal circuits. These data provide insights into the mechanisms underlying DEE-13, where seizures and intellectual disability are thought to be causally independent [12].

A key finding from our study is that hippocampal assembloids are capable of generating some of the canonical electrodynamic features of in vivo hippocampus. One of the most striking results that arose from this was the disordered theta-gamma coupling observed in the Hc+GE assembloids harboring *p*.*SCN8A* variants. Unlike the hyperexcitability seen in Mut Cx+GE assembloids, Mut Hc+GE assembloids displayed a loss of monophasic theta-gamma coupling, a pattern we also confirmed in vivo in human epilepsy patients. This aberrant coupling in the hippocampal models underscores the potential of these variants to disrupt hippocampal function, possibly contributing to the cognitive deficits observed in DEE-13 independent of seizures. This highlights the value of examining neural dynamics within specific brain circuits to understand how genetic variants contribute to diverse neurological and cognitive symptoms.

We used these findings to implement a predictive computational model of hippocampal function and driver of network dysfunction. The in silico data suggested that altered excitation/inhibition balance, resulting from decreased slow-spiking interneurons and increased excitatory neurons, may partly drive the network changes observed. Further immunohistochemical and snRNA sequencing analyses supported this prediction and expanded on the underlying changes in circuit development and composition from *p*.*SCN8A* expression. Notably, the selective loss of Reelin-expressing inhibitory interneurons, including O-LM and trilaminar neurons, in Hc+GE Mut assembloids highlights a specific vulnerability within the hippocampal circuit in DEE-13. These cells are associated with hippocampal oscillatory activity [49, 51, 52], and their loss likely contributes to disrupted theta-gamma coupling and high gamma/ripple activity, respectively, in Hc+GE assembloids with *p*.*SCN8A* mutations. These outcomes are indeed accounted for in our computational model by the incomplete restoration of monophasic theta-gamma coupling despite renormalizing the persistent sodium current (Extended Data Fig. 3a).

The value of this approach lies in transforming hiPSCs into in vitro structures capable of generating complex electrophysiological network dynamics, mimicking known in vivo human neural network signatures. This allows for the precise localization and analysis of specific abnormalities. Different brain regions in vivo, such as the cortex and hippocampus, do not form in isolation. Rather, they depend on cellular and neural network interactions that drive each region’s development and functional maturation. A better understanding of these interactions can come from complimentary in vivo studies of development in animal models, as well as future work utilizing more complex assembloid models. Nevertheless, studying these circuits independently provides important insights into specific abnormalities.

Looking forward, we propose further exploration of the observed network differences in Mut Cx+GE versus Mut Hc+GE. Our data suggest a permissive niche for developing a unique interneuron population in iCtrl Hc+GE (RELN+ SST and TL) absent in Cx+GE and inhibited in Hc+GE by gain-of-function *p*.*SCN8A* variants. Expression differences in excitatory genes (Extended Data Fig. 8) may play a role in establishing these region- and condition-specific developmental fates. A recent study [53] demonstrated that the presence of dysfunctional excitatory neurons can induce interneurons to change their relative abundance and identity, including a specific loss of O-LM defining characteristics, likely through aberrant maturation. This idea warrants further exploration with targeted depletion of *p*.*SCN8A* or specific blockers of NaV1.6 sodium channels to determine if these manipulations could hippocampal cellular changes and network dysfunction.

Leveraging the scalability of organoids, high-throughput screening of molecules that can reverse aberrant phenotypes in Cx+GE and Hc+GE assembloids could generate leads for clinical research to ameliorate DEE-13 symptoms. Such screening could identify novel compounds with potential therapeutic benefits for DEE-13 and other neurological conditions with similar pathophysiological mechanisms.

In summary, generating and analyzing distinct region-specific brain assembloids reveals divergent network dynamics and developmental trajectories in these brain regions with and without *p*.*SCN8A* variants. These models are powerful tools for dissecting the roles of specific neural circuit dynamics in health and disease and underscore the potential of regional assembloids to advance our understanding of complex neurological diseases. The insights gained hold promise for developing targeted and effective therapeutic interventions for conditions like DEE-13, where traditional approaches have limitations.

## 4 Acknowledgements

This work was supported the National Institutes of Health (#5K08NS119747 for R.A.S., #R01MH130061 and #R01DA051897 for B.G.N.), the Citizens United for Research in Epilepsy (#20204012 for R.A.S.), the Simons Foundation (#717153 for R.A.S.), and the UCLA Intellectual and Developmental Disabilities Research Center (#P50HD103557 for R.A.S. and B.G.N.).

## 6 Methods

### hiPSC culture and organoid generation

Human induced pluripotent stem cells (hiPSCs), including *p*.*SCN8A* mutant lines and controls, were generated by the Parent lab at the University of Michigan as previously described [15]. These hiPSCs, derived from de-identified patients reprogrammed from fibroblasts, were cultured using protocols employing pCXLE-hOCT3/4-shp53RNA, pCXLE-hUL, and pCXLE-hSK. The iCtrl for P1 was also generated and validated by Dr. Parent, confirmatory Sanger Sequencing is included here (Extended Data Fig. 9). All hiPSC experiments received prior approval from the University of California, Los Angeles (UCLA) Embryonic Stem Cell Research Oversight (ESCRO) committee and the Institutional Review Board.

Organoids representing the cortex (Cx), hippocampus (Hc), and ganglionic eminence (GE) were developed using cells from a female patient with the R1872*>*L mutation and a female patient with the V1592*>*L mutation, along with a CRISPR-corrected isogenic control for R1872*>*L. Organoids were induced toward a hippocampal fate by applying BMP4 along with CHIR 99021, a GSK3 inhibitor that acts as a Wnt agonist [9]. A GE (ganglionic eminence) fate was achieved by using agonists of the Sonic hedgehog (Shh) pathway [20], while a cortical fate was established by omitting these specific patterning factors [20]. Assembloids were created by fusing organoids of different types (Hc+GE and Cx+GE) using a modified version of the fusion protocol described in [4]. On day 56, organoids were bisected and combined in microcentrifuge tubes containing 400 *µ*l of medium—N2B27 for Cx+GE and Neurobasal for Hc+GE. These tubes were then incubated in a hyperoxic environment containing 5% CO_2_ and 40% O_2_ for 72 hours. After incubation, fused assembloids were transferred to 24-well oxygen-permeable dishes (Lumox, Sarstedt) and maintained under similar conditions with media changes every other day. Neuronal migration experiments involved the infection of individual organoids with 5 *µ*l of approximately 1.98 × 10^13^ ml^−1^ AAV1-tdTomato (pENN.AAV.CAG.tdTomato.WPRE.SV40, provided by J. M. Wilson, University of Pennsylvania Vector Core AV-1-PV3365) on day 56. Fusion of the infected organoids was executed three days post-infection, as previously described [4]. All experimental replicates represent assembloids derived from independent iPSC differentiations (referred to as “independently generated” above).

### Immunohistochemistry

Assembloids were immersion fixed in 4% paraformaldehyde, cryoprotected in 30% sucrose, frozen in Tissue-Tek Optimal Cutting Temperature medium (Sakura) and cryosectioned. Immunostaining was performed using previously published laboratory protocols [20]. Primary antibody staining was conducting using the following: rabbit anti-CAMK2 (ProteinTech, 13730-1-AP), 1:5,000 dilution; rat anti-CTIP2 (BCL11B; Abcam, ab18465), 1:1,000 dilution; rabbit anti-DLX1 (generous gift of S. K. Lee and J. Lee [54]), 1:3,000 dilution; guinea pig anti-DLX2 (generous gift of K. Yoshikawa and H. Shinagawa [55]), 1:3,000 dilution; mouse anti-GAD65 (BD Biosciences, 559931), 1:200 dilution; rabbit anti-GRIK4 (Invitrogen, PA5-111717), 1:100 dilution; mouse anti-NKX2-1 (Novocastra NCL-L-TTR-1), 1:500 dilution; goat anti-NRP2 (R&D Systems, AF2215), 1:40 dilution; rabbit anti-OLIG2 (EMD Millipore, AB9610), 1:5,000 dilution; rabbit anti-PAX6 (MBL International, PD022), 1:1,000 dilution; mouse anti-PROX1 (EMD Millipore, MAB5654), 1:200 dilution; rabbit anti-PV (Abcam, ab11427), 1:500 dilution; mouse anti-REELIN (MBL International, D3513), 1:300 dilution; mouse anti-SATB2 (Abcam, ab51502), 1:100 dilution; rat anti-SST (EMD Millipore MAB354), 1:100 dilution; rabbit anti-TBR1 (Abcam, ab31940), 1:2,000 dilution; chicken anti-TBR2 (EOMES; EMD Millipore, AB15894), 1:1,000 dilution; rabbit anti-ZBTB20 (Sigma-Aldrich, HPA016815), 1:100 dilution. Secondary antibodies used included the following conjugated donkey anti-species-specific IgG antibodies (Jackson ImmunoResearch): Alexa 488-, Cy3-, and Alexa 594-at a 1:1,000 dilution, and Alexa 647-at a 1:800 dilution. Hoechst 33342 was included in the secondary antibody mixture at a concentration of 1 *µ*g/ml to stain nuclei.

### Assembloid calcium imaging

The genetically encoded calcium indicator GCaMP6f was introduced into assembloids during days 88 to 95 through viral transduction using 5 *µ*L of 1.98 × 10^13^ GC mL^−1^ pAAV1.Syn.GCaMP7f.WPRE.SV40 virus [56], generously provided by D. Kim and the GENIE Project (Addgene viral preparation no. 100837-AAV1). Imaging was conducted 14 to 16 days post-infection utilizing a Leica Stellaris two-photon microscope equipped with a Coherent Chameleon tunable laser. Calcium transients were captured at an excitation wavelength of 920 nm employing a × 25 0.8-NA water-immersion objective (Nikon) at a frame rate of 31 Hz, with a resolution of 512 × 512 pixels and a field of view of 0.5 × 0.5 mm. Recordings were carried out in artificial cerebrospinal fluid (aCSF) supplemented with 10 mM HEPES to maintain a pH between 7.3 and 7.4, in the absence of *O*_2_*/CO*_2_ perfusion (refer to ‘Local field potential recordings’ section for further details).

Single-neuron calcium imaging data were extracted using a custom MATLAB script that leverages the CaImAn toolbox [57], specifically employing the CNMF (Constrained Nonnegative Matrix Factorization) algorithm [58] for source extraction and spike inference. The data underwent motion correction using the NoRMCorre package [59]. Components were initialized and iteratively updated, with relevant components selected through correlation and a pre-trained convolutional neural network classifier. The final processed data consisted of ΔF/F traces and inferred raster plots of action potentials.

### Assembloid local field potential recordings

Organoid recordings were conducted approximately between days 115 and 125. Live organoids were perfused with 250 nM kainic acid in artificial cerebrospinal fluid (aCSF), which consisted of 126 mM NaCl, 10 mM D-glucose, 1.2 mM MgCl_2_, 2 mM CaCl_2_, 5 mM KCl, 1.25 mM NaH_2_PO_4_, 1.5 mM sodium pyruvate, 1 mM L-glutamine, and 26 mM NaHCO_3_, to stimulate oscillatory network activities. The pH was maintained at 7.3–7.4, and aCSF was bubbled with 95% O_2_ and 5% CO_2_. Local field potential (LFP) activity was captured using a patch pipette filled with aCSF, connected to a field amplifier (A-M Systems, 3000). The signal was bandpass filtered between 0.1 and 1,000 Hz through an instrumentation amplifier (Brownlee BP Precision, 210A). Field potentials were digitized at 4,096 Hz using a National Instruments A/D board, interfaced with EVAN, a custom-designed LabView-based software from Thotec. Data analysis was performed using custom procedures developed in Matlab or Igor Pro.

### Human patient local field potential recordings

We reanalyzed previously published data [60] on hippocampal local field potentials from eight patients (four female, four male, ages 31-50, Extended Data Table 1) who underwent stereotactic implantation of bilateral intracranial depth electrodes (Integra or Ad-Tech, 5 mm spacing) at the University of California, Irvine Medical Center to identify seizure onset zones for potential surgical intervention. The study was approved by the institutional review boards at the University of California, Berkeley, and Irvine, with all subjects providing written informed consent. Electrode placements were based purely on clinical needs with MRI verification in the hippocampus. Electrode locations were confirmed using co-registered pre- and post-implantation T1-weighted MRI scans, with a high-resolution anatomical template (0.55 mm) aligned to each individual’s scan for precise localization. Only hippocampal recordings, specifically from DG/CA3 and CA1, were analyzed. Data were acquired with a Nihon Kohden system, filtered above 0.01 Hz, sampled at 5000 Hz, and analyzed using MATLAB with open-source toolboxes and custom scripts. During the initial study, participants viewed silent movie clips; however, here, we focused on task-free and seizure-free baseline hippocampal field potentials, examining theta-gamma phase-amplitude coupling monophasicity in each patient’s healthy and epileptic hippocampus.

### Theta-gamma coupling monophasicity

To analyze the degree of monophasic coupling between theta phase and gamma amplitude in our hippocampal assembloids, human epilepsy patients, and hippocampal circuit simulations, we utilized a custom MATLAB function. This function first defines a set of amplitude frequencies ranging from 25 to 115 Hz and applies a band-pass filter to the local field potential at each frequency band, followed by a Hilbert transform to extract the amplitude envelope. Simultaneously, the data is filtered within the theta band (3-10 Hz) and transformed to obtain the instantaneous phase. The relationship between the amplitude of gamma frequencies and the phase of theta is then computed by averaging the normalized amplitude of gamma at each theta phase bin, divided into 18 bins spanning from −*π* to *π*. This results in a radian modulogram matrix, where each row represents a gamma frequency and each column a theta phase bin. To quantify monophasicity, which indicates whether the amplitude of all gamma frequencies couples predominantly to one phase of theta, the matrix is correlated across all frequency pairs, and the median of these correlation coefficients (excluding the diagonal) is calculated as the monophasicity for each 10-second trial. This process is repeated for all trials in a given dataset, and the overall monophasicity for the organoid or patient is then taken as the median of these values.

### Instability analysis

To assess the instability of theta rhythms and single neuron spiking dynamics in our organoid models, human data, and simulations, we employed the modified 0-1 chaos test [61–63] alongside the estimation of the largest Lyapunov exponent for the simulations. The modified 0-1 chaos test involves driving a two-dimensional dynamical system with the recorded time-series data ϕ:

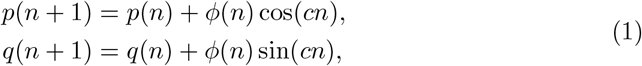

where *c* is a randomly chosen constant between 0 and 2*π*. The system’s trajectory, through the computation of the growth rate of the time-averaged mean square displacement of *p* and *q*, distinguishes chaotic from periodic dynamics. The mean square displacement is calculated as:

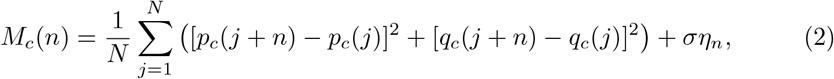

where *η*_*n*_ is a noise term. We set *σ* = 0.5, based on our prior work [42]. The correlation coefficient *K*_*c*_ = corr(*n, M*_*c*_(*n*)) is calculated for 100 unique values of *c*, and the median *K* value is used as the final statistic, with values approaching 1 indicating chaos and values near 0 suggesting periodicity. To estimate the instability of theta waves, the modified 0-1 chaos test was applied to local field potentials (real and simulated) bandpass filtered between 3-10 Hz; for estimating the instability of single-neuron spiking, we applied the test directly to the assembloid raster plots (inferred from the calcium imaging data) and to the raster plots of our simulated hippocampal circuit.

For our simulations, we estimated the largest Lyapunov exponent (which can only be estimated for in silico models) to directly quantify chaos. The simulations were run twice with a small perturbation in the initial membrane potentials of the neurons in the simulated hippocampal circuit, and the divergence *ϵ*(*t*) between these two runs was calculated. The stochastic largest Lyapunov exponent was then estimated from the divergence *ϵ*(*t*) between the initial and perturbed runs of the simulation, denoted 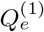 and 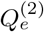, respectively, was calculated using the cumulative squared differences between their outputs:

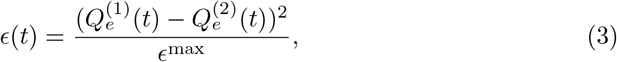

where *ϵ*^max^ represents the maximum divergence between the two simulation traces:

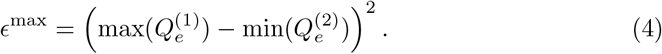

To estimate the largest Lyapunov exponent Λ, we calculate how the divergence *ϵ*(*t*) evolves over time:

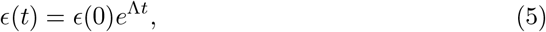

where *ϵ*(0) is the initial divergence at *t* = 0. The Lyapunov exponent Λ can then be inferred from the slope of the natural logarithm of *ϵ*(*t*) plotted against time *t*. Importantly, both runs, 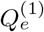 and 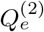, were subjected to identical noise conditions to ensure that any observed divergence is due to dynamical differences rather than stochastic variability. The computed slope provides a measure of the stochastic largest Lyapunov exponent, where a positive slope (and thus a positive stochastic largest Lypuanov exponent) indicates chaos/dynamical instability.

### Sample preparation for single-nuclei RNA sequencing

For each assembloid condition from the R1872*>*L *p*.*SCN8A* (iCtrl Cx+GE day 120, Mut P1 Cx+GE day 120, iCtrl Hc+GE day 120, Mut P1 Hc+GE day 120, iCtrl Hc+GE day 84, Mut P1 Hc+GE day 84), three batches of assembloids, each from a different differentiation, were pooled together into a single tube to be dissociated. In this manner, variability across differentiations was accounted for. Overall, between 3-11 assembloids per condition (across all batches) underwent the dissociation/sample preparation steps. All sample preparation steps were performed on dry ice or ice to minimize degradation effects from sample thawing or humidity. Tissue chunks were transferred to pre-cooled Dounce tissue grinder (Millipore Sigma, #D9063), and homogenized in RNase-free conditions with a light detergent (0.001% Triton100X/1%BSA/RNase Inhibitor (0.2U/µL)/0.5X Protease Inhibitor/1µM DTT/PBS). The homogenate was filtered through a 40 µm micro-cell strainer (Diagnocine, #FNK-HT-AMS-04002) to remove large material and centrifuged at 1000g for 8 min at 4ºC to pellet away from debris. The nuclei pellet was washed two more times with PBS/1%BSA/RNase Inhibitor and spun down at 500g for 5 mins. The pellet was then suspended in wash buffer for quality assessment and concentration measurement. Nuclei quality was evaluated based on shape, membrane integrity and chromatin structure under 40X magnification, and counted using a countess II machine. The isolated nuclei were then sent to the UCLA Technology Center for Genomics & Bioinformatics to construct expression libraries using the 10x Genomics Chromium Single Cell 3’ (v3 chemistry) system. The libraries aimed to recover 10K nuclei using a recommended concentration range. RNA sequencing on the NovaSeqS4 Illumina achieved an acceptable read depth for each condition (between 27,541-78,121 mean read pairs/cell for 4,613-9,813 cells per condition). All samples achieved *>* 96% genes mapped to the genome.

### Single-nuclei RNA sequencing analysis

The FASTQ file sequencing data for each assembloid condition (iCtrl Cx+GE day 120, Mut P1 Cx+GE day 120, iCtrl Hc+GE day 120, Mut P1 Hc+GE day 120, iCtrl Hc+GE day 84, Mut P1 Hc+GE day 84) were processed using the Cell Ranger 7.1.0 pipeline to generate count data. These were subsequently processed using custom Python v3.11.8 based scripts that implemented the Scanpy v1.9.8 single cell analysis package [64]. Briefly, genes that were expressed in less than 30 cells across all conditions were filtered out. Next, thresholds for total RNA counts and unique feature counts were determined for each condition via manual inspection in order to filter out multiplets and low quality cells. Manual thresholds were used as these counts were not normally or symmetrically distributed. Total RNA count thresholds typically ranged between 1000-15000 counts per cell, with 1000-8000 unique features per cell. Cells with *>* 1% mitochondrial RNA were also filtered out. Scrublet [65] was used to ensure adequate removal of doublets. Next, cell cycle analysis of S phase and G2/M phase gene markers (such as *MCM5, PCNA*, and *CDK1, NUSAP1*, respectively) was performed to confirm that relatively few S and G2/M phase cells were present in the data. After these initial filtering steps, the remaining high-quality cells were integrated across all 6 conditions using the single cell variational inference package scVI v1.1.2 [66] (using 2 hidden layers with 128 nodes per layer, a latent space of 30 dimensions, and gene likelihood modeled by a zero-inflated negative binomial distribution). Nearest neighbors were calculated using a local neighborhood size of 15 with distance metric based on the Pearson correlation coefficient given previously reported optimal performance [67] prior to UMAP projection. Visual inspection was subsequently performed to verify reasonable integration across batches. Leiden clustering using multiple resolutions in a highly iterative manner was then performed for gross cell type labeling. ExN and IINs were subsequently isolated and each was independently reintegrated across batches using the prior method to perform subtype clustering and labeling. ExN and IN cluster labeling (Extended Data Fig. 5 and 6, respectively) utilized previously reported canonical expression markers [68–75].

After cell labeling, a scANVI integration step [76] was utilized to generate the UMAP plots in Fig. 1, Fig. 6, and Extended Data Fig. 7 in order to maximize biological conservation during batch integration. DGE analysis was performed using Scanpy’s built-in functions (Wilcoxon method with tie-correct and Bonferroni correction) given their previously reported optimal performance [77, 78]. DGE significance testing utilized a threshold of *p* − *adjusted* < 0.05 and log-fold change of *>* 0.5 and was performed individually for each cell type (where at least 30 cells were present across all conditions) to focus the DGE analysis on within-cell-type differences (as cell-type differences were analyzed independently). To check for pathological changes through DGE analysis, significantly downregulated genes were compared to the list of genes that, when dysfunctional, are associated with epilepsy (CUI: C0014544) and intellectual disability (CUI: C3714756) using DisGenNet (https://www.disgenet.org/search) [50]. Significantly upregulated genes from DGE analysis were also compared to the list of genes associated with glutamatergic/excitatory functions. Gene ontology analysis was performed for all upregulated and downregulated genes using the Gene Set Enrichment Analysis package in Python (GSEApy) [79] with the human *GO Biological Process 2021* dataset and a significance cutoff of *p*− *adjusted* < 0.05. Super-clustering was performed using the GO-Figure package [80]. For the excitatory gene analysis in Extended Data Fig. 8, 66 unique genes from the union of GO:0098976 (excitatory chemical synaptic transmission), GO:0060079 (excitatory postsynaptic potential), GO:2000463 (positive regulation of excitatory postsynaptic potential), GO:0035249 (synaptic transmission, glutamatergic), and GO:0051968 (positive regulation of synaptic transmission, glutamatergic) were used that best represented the set of excitatory genes. This analysis was performed across the different conditions and was not stratified by cell type; a significance threshold of *p* − *adjusted* < 0.05 and a log-fold change threshold *>* 0.5 were used.

### Hippocampal circuit simulation

The O-LM cell model is adapted from the multicomponent model described in Saraga et al. [81], which was initially developed as a multicompartment model. This model was later simplified into a single-compartment model by Tort et al [44]. Building on this simplified model, we have introduced a persistent sodium current (NaP), enhancing its ability to model the pathophysiology of DEE-13.

The O-LM cell model is described by the current-balance equation:

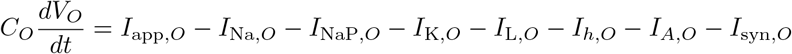

with *V*_*O*_ as the membrane potential, *C*_*O*_ = 1.3*µF/cm*^2^ as the capacitance, *I*_app,*O*_ the applied current, and *I*_syn,*O*_ the synaptic current. The leak current *I*_*L,O*_ has a conductance of 0.05*mS/cm*^2^ and a reversal potential of − 70*mV*. The currents in this equation are defined as:

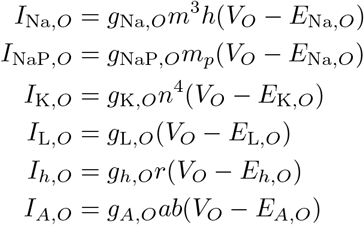

where the gating variables are governed by:

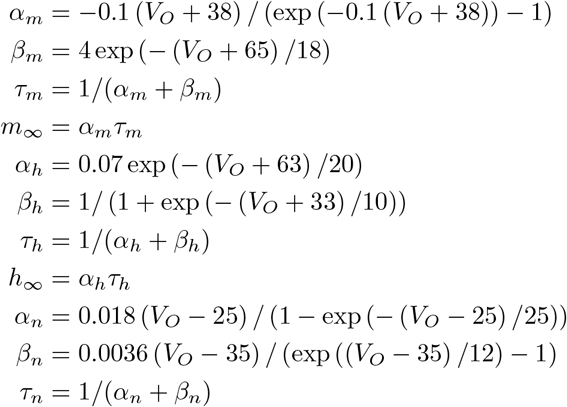

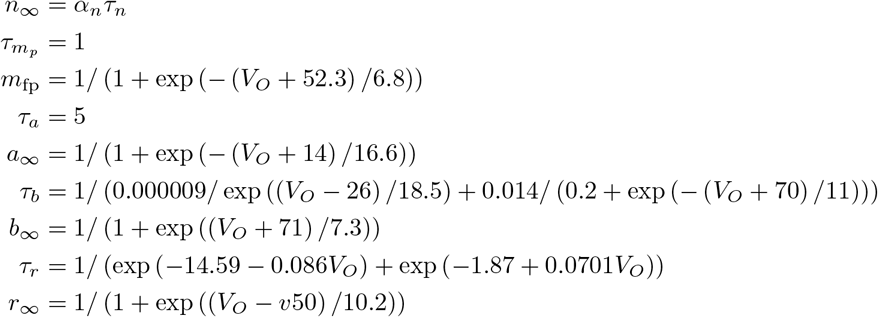

The basket cell model, adapting the characteristics of a fast-spiking interneuron as initially described by Wang and Buzsaki [82] and later refined by Tort et al. [44], likewise incorporates a persistent sodium current. It is a single-compartment model described by:

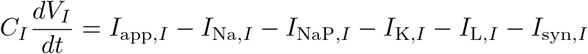

where *V*_*I*_ denotes the membrane potential, *C*_*I*_ = 1*µF/cm*^2^ indicates the membrane capacitance, *I*_app,*I*_ is the applied current, *I*_syn,*I*_ is the total synaptic current, and *I*_*L,I*_ = *g*_*L,I*_ (*V*_*I*_ − *E*_*L,I*_) with a conductance of 0.1*mS/cm*^2^ and a reversal potential of −65*mV*. All currents are measured in *µA/cm*^2^. The currents in this equation are defined as:

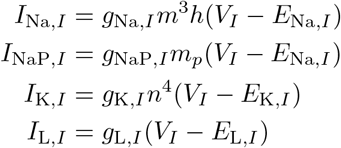

where gating variables and their kinetics are given by:

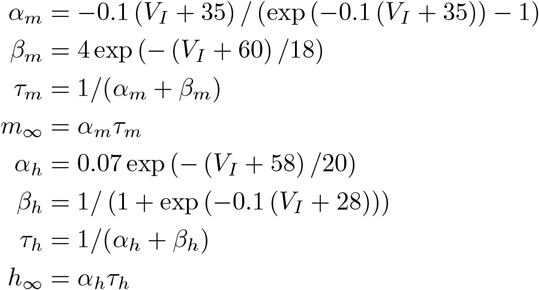

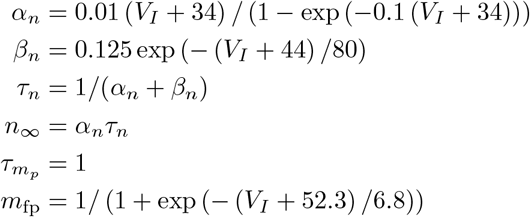

Finally, the pyramidal cell model is based on the multicompartmental framework first described by Migliore et al. [83], which was subsequently adapted into a more computationally efficient model by Tort et al. [44]. In our current work, we have added a persistent sodium current (NaP) to this model, reflecting a deeper understanding of the pyramidal neurons’ intrinsic properties that affect their long-term excitability and connectivity within neural circuits.

The pyramidal cell model includes basal dendrites (200 *µm* length, 2 *µm* diameter), a soma (20 *µm* length, 20 *µm* diameter), andt hree apical dendrites (each 150 *µm* length, 2 *µm* diameter). Cytoplasmatic resistivity (*Ra*) is set at 150 Ω · *cm*. The model for each compartment *k* is governed by:

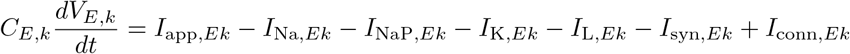

where the currents are defined as:

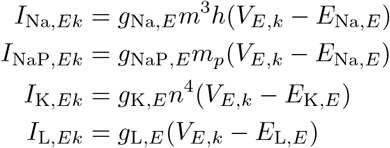

and the gating variables and their kinetics are given by:

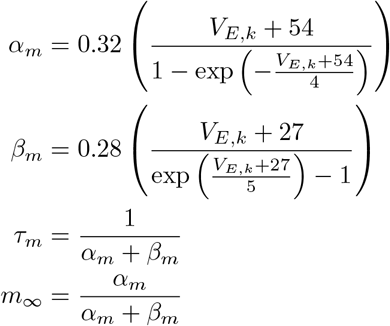

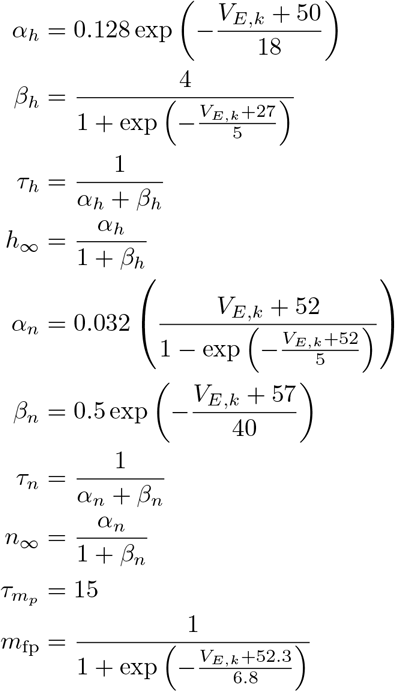

To simulate the electrophysiological characteristics of iCtrl Hc+GE assembloids, specifically recreating the observed monophasic theta-gamma coupling, we utilized a genetic optimization approach. This process involved adjusting the parameters of the above-described hippocampal neural network model incorporating O-LM cells, basket cells, and pyramidal cells. The genetic algorithm optimized several key parameters to achieve a close match with the observed electrophysiological data. These parameters included the strength and noise level of the driving currents applied to these cells (measured in picoamperes, pA), the strengths of transient sodium, persistent sodium, and potassium currents across all cell types, and the connectivity between all pairs of cell types (e.g., O-LM to O-LM, O-LM to pyramidal, etc.). To simulate Mut Hc+GE assembloids, we again employed a genetic algorithm, with the iCtrl simulation parameters as the starting point, and optimized the cell numbers and strength of persistent sodium currents to recreate the aberrant theta rhythmicity and theta-gamma coupling, but preserved theta power, of the Mut Hc+GE assembloids.

### Statistical Information

Graph generation and statistical analyses were performed either on GraphPad Prism 10 or Matlab software. In cases where there were repeated measurements within a single sample (e.g. repeated calcium indicator measurements from the same organoid) a linear mixed effects model was used to determine statistical differences. All other samples were subjected to Shapiro–Wilk and Kolmogorov–Smirnov normality testing.

Non-normal samples were analyzed by a one- or two-tailed Mann–Whitney U test or Kruskal–Wallis test followed by a Dunn’s multiple-comparison test. Normally distributed data were analyzed by a two-tailed Student’s t-test or ANOVA with post hoc Tukey’s multiple-comparison test unless specified in the figure legend. Sample sizes were not pre-determined but are comparable to prior publications. Organoids were generated from hiPSCs in batches of 96 organoids per plating and subjected to visual inspection with each media change (q2-3 days) and more detailed visual quality control checks at the following timepoints: day 6, 18, and 35. Of the samples that passed these quality control checks, individual organoids were randomly selected for experiments. All cell counting was performed by an individual blinded to experimental group and data were unblinded only after all counts were completed for a specific immunostain. Other data collection and analysis were not performed blinded. No data points were excluded from analysis for any reason. Bar graphs indicate the mean and standard deviation of the data, and all individual data points are plotted to demonstrate the full distribution of values. Figure legends specify sample numbers and exact *p* values for all statistical tests. Each n value represents an independent experiment performed on an assembloid or organoid generated from an independent hiPSC differentiation.

## 7 Data Availability

Raw and processed snRNAseq data will be deposited in the NCBI Gene Expression Omnibus and the raw gene expression analysis results will be made publicly available once accepted for publication. The authors declare that all other data supporting the findings of this study are available within the main text of the paper and the Extended Data and Supplementary Data files.

## 8 Code Availability

All code used in data analysis was previously generated and published by others. Any modifications have been described in the Methods section.

## References

[1] Di Lullo, E., Kriegstein, A.R.: The use of brain organoids to investigate neural development and disease. Nature Reviews Neuroscience 18(10), 573–584 (2017)

[2] Amin, N.D., Paşca, S.P.: Building models of brain disorders with three-dimensional organoids. Neuron 100(2), 389–405 (2018)

[3] Qian, X., Song, H., Ming, G.-l.: Brain organoids: advances, applications and challenges. Development 146(8), 166074 (2019)

[4] Samarasinghe, R.A., Miranda, O.A., Buth, J.E., Mitchell, S., Ferando, I., Watanabe, M., Allison, T.F., Kurdian, A., Fotion, N.N., Gandal, M.J., et al.: Identification of neural oscillations and epileptiform changes in human brain organoids. Nature neuroscience 24(10), 1488–1500 (2021)

[5] Trujillo, C.A., Gao, R., Negraes, P.D., Gu, J., Buchanan, J., Preissl, S., Wang, A., Wu, W., Haddad, G.G., Chaim, I.A., et al.: Complex oscillatory waves emerging from cortical organoids model early human brain network development. Cell stem cell 25(4), 558–569 (2019)

[6] Sharf, T., Molen, T., Glasauer, S.M., Guzman, E., Buccino, A.P., Luna, G., Cheng, Z., Audouard, M., Ranasinghe, K.G., Kudo, K., et al.: Functional neuronal circuitry and oscillatory dynamics in human brain organoids. Nature communications 13(1), 4403 (2022)

[7] Kadoshima, T., Sakaguchi, H., Nakano, T., Soen, M., Ando, S., Eiraku, M., Sasai, Y.: Self-organization of axial polarity, inside-out layer pattern, and species-specific progenitor dynamics in human es cell–derived neocortex. Proceedings of the National Academy of Sciences 110(50), 20284–20289 (2013)

[8] Xiang, Y., Tanaka, Y., Cakir, B., Patterson, B., Kim, K.-Y., Sun, P., Kang, Y.-J., Zhong, M., Liu, X., Patra, P., et al.: hesc-derived thalamic organoids form reciprocal projections when fused with cortical organoids. Cell stem cell 24(3), 487–497 (2019)

[9] Sakaguchi, H., Kadoshima, T., Soen, M., Narii, N., Ishida, Y., Ohgushi, M., Takahashi, J., Eiraku, M., Sasai, Y.: Generation of functional hippocampal neurons from self-organizing human embryonic stem cell-derived dorsomedial telencephalic tissue. Nature communications 6(1), 1–11 (2015)

[10] Birey, F., Andersen, J., Makinson, C.D., Islam, S., Wei, W., Huber, N., Fan, H.C., Metzler, K.R.C., Panagiotakos, G., Thom, N., et al.: Assembly of functionally integrated human forebrain spheroids. Nature 545(7652), 54–59 (2017)

[11] Miura, Y., Li, M.-Y., Birey, F., Ikeda, K., Revah, O., Thete, M.V., Park, J.-Y., Puno, A., Lee, S.H., Porteus, M.H., et al.: Generation of human striatal organoids and cortico-striatal assembloids from human pluripotent stem cells. Nature Biotechnology 38(12), 1421–1430 (2020)

[12] Specchio, N., Curatolo, P.: Developmental and epileptic encephalopathies: what we do and do not know. Brain 144(1), 32–43 (2021)

[13] Johannesen, K.M., Liu, Y., Koko, M., Gjerulfsen, C.E., Sonnenberg, L., Schubert, J., Fenger, C.D., Eltokhi, A., Rannap, M., Koch, N.A., et al.: Genotype-phenotype correlations in scn8a-related disorders reveal prognostic and therapeutic implications. Brain 145(9), 2991–3009 (2022)

[14] Lopez-Santiago, L.F., Yuan, Y., Wagnon, J.L., Hull, J.M., Frasier, C.R., O’Malley, H.A., Meisler, M.H., Isom, L.L.: Neuronal hyperexcitability in a mouse model of scn8a epileptic encephalopathy. Proceedings of the National Academy of Sciences 114(9), 2383–2388 (2017)

[15] Tidball, A.M., Lopez-Santiago, L.F., Yuan, Y., Glenn, T.W., Margolis, J.L., Clayton Walker, J., Kilbane, E.G., Miller, C.A., Martina Bebin, E., Scott Perry, M., et al.: Variant-specific changes in persistent or resurgent sodium current in scn8a-related epilepsy patient-derived neurons. Brain 143(10), 3025–3040 (2020)

[16] Liu, Y., Schubert, J., Sonnenberg, L., Helbig, K.L., Hoei-Hansen, C.E., Koko, M., Rannap, M., Lauxmann, S., Huq, M., Schneider, M.C., et al.: Neuronal mechanisms of mutations in scn8a causing epilepsy or intellectual disability. Brain 142(2), 376–390 (2019)

[17] Veeramah, K.R., O’Brien, J.E., Meisler, M.H., Cheng, X., Dib-Hajj, S.D., Waxman, S.G., Talwar, D., Girirajan, S., Eichler, E.E., Restifo, L.L., et al.: De novo pathogenic scn8a mutation identified by whole-genome sequencing of a family quartet affected by infantile epileptic encephalopathy and sudep. The American Journal of Human Genetics 90(3), 502–510 (2012)

[18] Wagnon, J.L., Barker, B.S., Hounshell, J.A., Haaxma, C.A., Shealy, A., Moss, T., Parikh, S., Messer, R.D., Patel, M.K., Meisler, M.H.: Pathogenic mechanism of recurrent mutations of scn8a in epileptic encephalopathy. Annals of clinical and translational neurology 3(2), 114–123 (2016)

[19] Ottolini, M., Barker, B.S., Gaykema, R.P., Meisler, M.H., Patel, M.K.: Aberrant sodium channel currents and hyperexcitability of medial entorhinal cortex neurons in a mouse model of scn8a encephalopathy. Journal of Neuroscience 37(32), 7643–7655 (2017)

[20] Watanabe, M., Buth, J.E., Vishlaghi, N., Torre-Ubieta, L., Taxidis, J., Khakh, B.S., Coppola, G., Pearson, C.A., Yamauchi, K., Gong, D., et al.: Self-organized cerebral organoids with human-specific features predict effective drugs to combat zika virus infection. Cell reports 21(2), 517–532 (2017)

[21] Götz, M., Stoykova, A., Gruss, P.: Pax6 controls radial glia differentiation in the cerebral cortex. Neuron 21(5), 1031–1044 (1998)

[22] Bulfone, A., Martinez, S., Marigo, V., Campanella, M., Basile, A., Quaderi, N., Gattuso, C., Rubenstein, J.L., Ballabio, A.: Expression pattern of the tbr2 (eomesodermin) gene during mouse and chick brain development. Mechanisms of development 84(1-2), 133–138 (1999)

[23] Arlotta, P., Molyneaux, B.J., Chen, J., Inoue, J., Kominami, R., Macklis, J.D.: Neuronal subtype-specific genes that control corticospinal motor neuron development in vivo. Neuron 45(2), 207–221 (2005)

[24] Britanova, O., Juan Romero, C., Cheung, A., Kwan, K.Y., Schwark, M., Gyorgy, A., Vogel, T., Akopov, S., Mitkovski, M., Agoston, D., et al.: Satb2 is a postmitotic determinant for upper-layer neuron specification in the neocortex. Neuron 57(3), 378–392 (2008)

[25] Elias, L.A., Potter, G.B., Kriegstein, A.R.: A time and a place for nkx2-1 in interneuron specification and migration. Neuron 59(5), 679–682 (2008)

[26] Petryniak, M.A., Potter, G.B., Rowitch, D.H., Rubenstein, J.L.: Dlx1 and dlx2 control neuronal versus oligodendroglial cell fate acquisition in the developing forebrain. Neuron 55(3), 417–433 (2007)

[27] Anderson, S.A., Qiu, M., Bulfone, A., Eisenstat, D.D., Meneses, J., Pedersen, R., Rubenstein, J.L.: Mutations of the homeobox genes dlx-1 and dlx-2 disrupt the striatal subventricular zone and differentiation of late born striatal neurons. Neuron 19(1), 27–37 (1997)

[28] Esclapez, M., Tillakaratne, N., Kaufman, D.L., Tobin, A.J., Houser, C.R.: Comparative localization of two forms of glutamic acid decarboxylase and their mrnas in rat brain supports the concept of functional differences between the forms. Journal of Neuroscience 14(3), 1834–1855 (1994)

[29] Chen, H., Chédotal, A., He, Z., Goodman, C.S., Tessier-Lavigne, M.: Neuropilin-2, a novel member of the neuropilin family, is a high affinity receptor for the semaphorins sema e and sema iv but not sema iii. Neuron 19(3), 547–559 (1997)

[30] Iwano, T., Masuda, A., Kiyonari, H., Enomoto, H., Matsuzaki, F.: Prox1 postmitotically defines dentate gyrus cells by specifying granule cell identity over ca3 pyramidal cell fate in the hippocampus. Development 139(16), 3051–3062 (2012)

[31] Grove, E.A., Tole, S.: Patterning events and specification signals in the developing hippocampus. Cerebral cortex 9(6), 551–561 (1999)

[32] Tole, S., Christian, C., Grove, E.A.: Early specification and autonomous development of cortical fields in the mouse hippocampus. Development 124(24), 4959–4970 (1997)

[33] Simon, R., Brylka, H., Schwegler, H., Venkataramanappa, S., Andratschke, J., Wiegreffe, C., Liu, P., Fuchs, E., Jenkins, N.A., Copeland, N.G., et al.: A dual function of bcl11b/ctip2 in hippocampal neurogenesis. The EMBO journal 31(13), 2922–2936 (2012)

[34] McEvilly, R.J., Diaz, M.O., Schonemann, M.D., Hooshmand, F., Rosenfeld, M.G.: Transcriptional regulation of cortical neuron migration by pou domain factors. Science 295(5559), 1528–1532 (2002)

[35] Sugitani, Y., Nakai, S., Minowa, O., Nishi, M., Jishage, K.-i., Kawano, H., Mori, K., Ogawa, M., Noda, T.: Brn-1 and brn-2 share crucial roles in the production and positioning of mouse neocortical neurons. Genes & development 16(14), 1760–1765 (2002)

[36] Eng, L., Vanderhaeghen, J., Bignami, A., Gerstl, B.: An acidic protein isolated from fibrous astrocytes. Brain research 28(2), 351–354 (1971)

[37] Terrigno, M., Busti, I., Alia, C., Pietrasanta, M., Arisi, I., D’Onofrio, M., Caleo, M., Cremisi, F.: Neurons generated by mouse escs with hippocampal or cortical identity display distinct projection patterns when co-transplanted in the adult brain. Stem cell reports 10(3), 1016–1029 (2018)

[38] Suresh, V., Muralidharan, B., Pradhan, S.J., Bose, M., D’Souza, L., Parichha, A., Reddy, P.C., Galande, S., Tole, S.: Regulation of chromatin accessibility and gene expression in the developing hippocampal primordium by lim-hd transcription factor lhx2. PLoS Genetics 19(8), 1010874 (2023)

[39] Buzsáki, G.: Hippocampal sharp wave-ripple: A cognitive biomarker for episodic memory and planning. Hippocampus 25(10), 1073–1188 (2015)

[40] Liu, A.A., Henin, S., Abbaspoor, S., Bragin, A., Buffalo, E.A., Farrell, J.S., Foster, D.J., Frank, L.M., Gedankien, T., Gotman, J., et al.: A consensus statement on detection of hippocampal sharp wave ripples and differentiation from other fast oscillations. Nature communications 13(1), 6000 (2022)

[41] Colgin, L.L.: Theta–gamma coupling in the entorhinal–hippocampal system. Current opinion in neurobiology 31, 45–50 (2015)

[42] Toker, D., Sommer, F.T., D’Esposito, M.: A simple method for detecting chaos in nature. Communications biology 3(1), 11 (2020)

[43] Gottwald, G.A., Melbourne, I.: The 0-1 test for chaos: A review. Chaos detection and predictability, 221–247 (2016)

[44] Tort, A.B., Rotstein, H.G., Dugladze, T., Gloveli, T., Kopell, N.J.: On the formation of gamma-coherent cell assemblies by oriens lacunosum-moleculare interneurons in the hippocampus. Proceedings of the National Academy of Sciences 104(33), 13490–13495 (2007)

[45] He, L., Caudill, M.S., Jing, J., Wang, W., Sun, Y., Tang, J., Jiang, X., Zoghbi, H.Y.: A weakened recurrent circuit in the hippocampus of rett syndrome mice disrupts long-term memory representations. Neuron 110(10), 1689–1699 (2022)

[46] Peng, Z., Zhang, N., Wei, W., Huang, C.S., Cetina, Y., Otis, T.S., Houser, C.R.: A reorganized gabaergic circuit in a model of epilepsy: evidence from optogenetic labeling and stimulation of somatostatin interneurons. Journal of Neuroscience 33(36), 14392–14405 (2013)

[47] Orban-Kis, K., Szabadi, T., Szilagyi, T.: The loss of ivy cells and the hippocampal input modulatory o-lm cells contribute to the emergence of hyperexcitability in the hippocampus. Rom J Morphol Embryol 56(1), 155–161 (2015)

[48] Wyeth, M., Buckmaster, P.S.: Lack of hyperinhibition of oriens lacunosum-moleculare cells by vasoactive intestinal peptide-expressing cells in a model of temporal lobe epilepsy. Eneuro 8(6) (2021)

[49] Ferraguti, F., Klausberger, T., Cobden, P., Baude, A., Roberts, J.D.B., Szucs, P., Kinoshita, A., Shigemoto, R., Somogyi, P., Dalezios, Y.: Metabotropic glutamate receptor 8-expressing nerve terminals target subsets of gabaergic neurons in the hippocampus. Journal of Neuroscience 25(45), 10520–10536 (2005)

[50] Piñero, J., Ramírez-Anguita, J.M., Saüch-Pitarch, J., Ronzano, F., Centeno, E., Sanz, F., Furlong, L.I.: The disgenet knowledge platform for disease genomics: 2019 update. Nucleic acids research 48(D1), 845–855 (2020)

[51] Jansen, N.A., Perez, C., Schenke, M., Beurden, A.W., Dehghani, A., Voskuyl, R.A., Thijs, R.D., Ullah, G., Maagdenberg, A.M., Tolner, E.A.: Impaired θ-γ coupling indicates inhibitory dysfunction and seizure risk in a dravet syndrome mouse model. Journal of Neuroscience 41(3), 524–537 (2021)

[52] Ponzi, A., Dura-Bernal, S., Migliore, M.: Theta-gamma phase amplitude coupling in a hippocampal ca1 microcircuit. PLOS Computational Biology 19(3), 1010942 (2023)

[53] Wu, S.J., Dai, M., Yang, S.-P., McCann, C., Qiu, Y., Marrero, G.J., Stogsdill, J.A., Di Bella, D.J., Xu, Q., Farhi, S.L., Macosko, E.Z., Chen, F., Fishell, G.: Pyramidal neurons proportionately alter the identity and survival of specific cortical interneuron subtypes. bioRxiv (2024)

[54] Lee, B., Kim, J., An, T., Kim, S., Patel, E.M., Raber, J., Lee, S.-K., Lee, S., Lee, J.W.: Dlx1/2 and otp coordinate the production of hypothalamic ghrh-and agrp-neurons. Nature communications 9(1), 2026 (2018)

[55] Kuwajima, T., Nishimura, I., Yoshikawa, K.: Necdin promotes gabaergic neuron differentiation in cooperation with dlx homeodomain proteins. Journal of Neuroscience 26(20), 5383–5392 (2006)

[56] Chen, T.-W., Wardill, T.J., Sun, Y., Pulver, S.R., Renninger, S.L., Baohan, A., Schreiter, E.R., Kerr, R.A., Orger, M.B., Jayaraman, V., et al.: Ultrasensitive fluorescent proteins for imaging neuronal activity. Nature 499(7458), 295–300 (2013)

[57] Giovannucci, A., Friedrich, J., Gunn, P., Kalfon, J., Brown, B.L., Koay, S.A., Taxidis, J., Najafi, F., Gauthier, J.L., Zhou, P., et al.: Caiman an open source tool for scalable calcium imaging data analysis. elife 8, 38173 (2019)

[58] Pnevmatikakis, E.A., Soudry, D., Gao, Y., Machado, T.A., Merel, J., Pfau, D., Reardon, T., Mu, Y., Lacefield, C., Yang, W., et al.: Simultaneous denoising, deconvolution, and demixing of calcium imaging data. Neuron 89(2), 285–299 (2016)

[59] Pnevmatikakis, E.A., Giovannucci, A.: Normcorre: An online algorithm for piece-wise rigid motion correction of calcium imaging data. Journal of neuroscience methods 291, 83–94 (2017)

[60] Zheng, J., Anderson, K.L., Leal, S.L., Shestyuk, A., Gulsen, G., Mnatsakanyan, L., Vadera, S., Hsu, F.P., Yassa, M.A., Knight, R.T., et al.: Amygdala-hippocampal dynamics during salient information processing. Nature communications 8(1), 14413 (2017)

[61] Gottwald, G.A., Melbourne, I.: A new test for chaos in deterministic systems. In: Proceedings of the Royal Society of London A: Mathematical, Physical and Engineering Sciences, vol. 460, pp. 603–611 (2004). The Royal Society

[62] Gottwald, G.A., Melbourne, I.: Testing for chaos in deterministic systems with noise. Physica D: Nonlinear Phenomena 212(1-2), 100–110 (2005)

[63] Dawes, J., Freeland, M.: The ‘0–1 test for chaos’ and strange nonchaotic attractors. preprint (2008)

[64] Wolf, F.A., Angerer, P., Theis, F.J.: Scanpy: large-scale single-cell gene expression data analysis. Genome biology 19, 1–5 (2018)

[65] Wolock, S.L., Lopez, R., Klein, A.M.: Scrublet: computational identification of cell doublets in single-cell transcriptomic data. Cell systems 8(4), 281–291 (2019)

[66] Lopez, R., Regier, J., Cole, M.B., Jordan, M.I., Yosef, N.: Deep generative modeling for single-cell transcriptomics. Nature methods 15(12), 1053–1058 (2018)

[67] Watson, E.R., Mora, A., Taherian Fard, A., Mar, J.C.: How does the structure of data impact cell–cell similarity? evaluating how structural properties influence the performance of proximity metrics in single cell rna-seq data. Briefings in Bioinformatics 23(6), 387 (2022)

[68] Cembrowski, M.S., Wang, L., Sugino, K., Shields, B.C., Spruston, N.: Hipposeq: a comprehensive rna-seq database of gene expression in hippocampal principal neurons. elife 5, 14997 (2016)

[69] Mukhtar, T., Taylor, V.: Untangling cortical complexity during development. Journal of experimental neuroscience 12, 1179069518759332 (2018)

[70] Matho, K.S., Huilgol, D., Galbavy, W., He, M., Kim, G., An, X., Lu, J., Wu, P., Di Bella, D.J., Shetty, A.S., et al.: Genetic dissection of the glutamatergic neuron system in cerebral cortex. Nature 598(7879), 182–187 (2021)

[71] Mayer, C., Hafemeister, C., Bandler, R.C., Machold, R., Batista Brito, R., Jaglin, X., Allaway, K., Butler, A., Fishell, G., Satija, R.: Developmental diversification of cortical inhibitory interneurons. Nature 555(7697), 457–462 (2018)

[72] Allison, T., Langerman, J., Sabri, S., Otero-Garcia, M., Lund, A., Huang, J., Wei, X., Samarasinghe, R.A., Polioudakis, D., Mody, I., et al.: Defining the nature of human pluripotent stem cell-derived interneurons via single-cell analysis. Stem cell reports 16(10), 2548–2564 (2021)

[73] Munguba, H., Chattopadhyaya, B., Nilsson, S., Carriço, J.N., Memic, F., Oberst, P., Batista-Brito, R., Muñoz-Manchado, A.B., Wegner, M., Fishell, G., et al.: Postnatal sox6 regulates synaptic function of cortical parvalbumin-expressing neurons. Journal of Neuroscience 41(43), 8876–8886 (2021)

[74] Harris, K.D., Hochgerner, H., Skene, N.G., Magno, L., Katona, L., Bengtsson Gonzales, C., Somogyi, P., Kessaris, N., Linnarsson, S., Hjerling-Leffler, J.: Classes and continua of hippocampal ca1 inhibitory neurons revealed by single-cell transcriptomics. PLoS biology 16(6), 2006387 (2018)

[75] Pavon, N., Diep, K., Yang, F., Sebastian, R., Martinez-Martin, B., Ranjan, R., Sun, Y., Pak, C.: Patterning ganglionic eminences in developing human brain organoids using a morphogen-gradient-inducing device. Cell Reports Methods 4(1) (2024)

[76] Xu, C., Lopez, R., Mehlman, E., Regier, J., Jordan, M.I., Yosef, N.: Probabilistic harmonization and annotation of single-cell transcriptomics data with deep generative models. Molecular systems biology 17(1), 9620 (2021)

[77] Li, Y., Ge, X., Peng, F., Li, W., Li, J.J.: Exaggerated false positives by popular differential expression methods when analyzing human population samples. Genome biology 23(1), 79 (2022)

[78] Pullin, J.M., McCarthy, D.J.: A comparison of marker gene selection methods for single-cell rna sequencing data. Genome Biology 25(1), 56 (2024)

[79] Fang, Z., Liu, X., Peltz, G.: Gseapy: a comprehensive package for performing gene set enrichment analysis in python. Bioinformatics 39(1), 757 (2023)

[80] Reijnders, M.J., Waterhouse, R.M.: Summary visualizations of gene ontology terms with go-figure! Frontiers in Bioinformatics 1, 638255 (2021)

[81] Saraga, F., Wu, C., Zhang, L., Skinner, F.: Active dendrites and spike propagation in multicompartment models of oriens-lacunosum/moleculare hippocampal interneurons. The Journal of physiology 552(3), 673–689 (2003)

[82] Wang, X.-J., Buzsáki, G.: Gamma oscillation by synaptic inhibition in a hippocampal interneuronal network model. Journal of neuroscience 16(20), 6402–6413 (1996)

[83] Migliore, M., Messineo, L., Ferrante, M.: Dendritic i h selectively blocks temporal summation of unsynchronized distal inputs in ca1 pyramidal neurons. Journal of computational neuroscience 16, 5–13 (2004)

