## Extended Data Figures and Tables for "Modeling Cortical Versus Hippocampal Network Dysfunction in a Human Brain Assembloid Model of Epilepsy and Intellectual Disability"

### Modeling Cortical Versus Hippocampal Network Dysfunction in a Human Brain Assembloid Model of Epilepsy and Intellectual Disability - Extended Data Figures and Tables

Colin M. McCrimmon & Daniel Toker[1†], Marie Pahos[1], Kevin Lozano[1], Jack J. Lin[2,3], Jack Parent[4,5,6], Andrew Tidball[5], Jie Zheng[7,8], László Molnár[9], Istvan Mody[1,10], Bennett G. Novitch[11,12,13], Ranmal A. Samarasinghe[1,11,12,13]

[1]Department of Neurology, University of California, Los Angeles, David Geffen School of Medicine, Los Angeles, 90095, CA, USA.
[2]Department of Neurology, University of California, Davis, School of Medicine, Sacramento, 95817, CA, USA.
[3]Center for Mind and Brain, University of California, Davis, Davis, 95618, CA, USA.
[4]Department of Neurology, University of Michigan School of Medicine, Ann Arbor, 48109, MI, USA.
[5]Michigan Neuroscience Institute, University of Michigan, Ann Arbor, 48109, MI, USA.
[6]VA Ann Arbor Healthcare System, Ann Arbor, 48105, MI, USA.
[7]Department of Biomedical Engineering, University of California, Davis, Davis, 95616, CA, USA.
[8]Department of Neurological Surgery, University of California, Davis, School of Medicine, Sacramento, 95817, CA, USA.
[9]Department of Electrical Engineering, Sapientia Hungarian University of Transylvania, Târgu-Mureş/Corunca, 540485, ROU.
[10]Department of Physiology, University of California, Los Angeles, David Geffen School of Medicine, Los Angeles, 90095, CA, USA.
[11]Department of Neurobiology, University of California, Los Angeles, Los Angeles, 90095, CA, USA.

[12]Eli and Edythe Broad Center for Regenerative Medicine and Stem Cell Research, University of California, Los Angeles, Los Angeles, 90095, CA, USA.
[13]Intellectual Development and Disabilities Research Center, University of California, Los Angeles, Los Angeles, 90095, CA, USA.

Contributing authors:;
†These authors contributed equally to this work.

### 1 Extended Data Figures

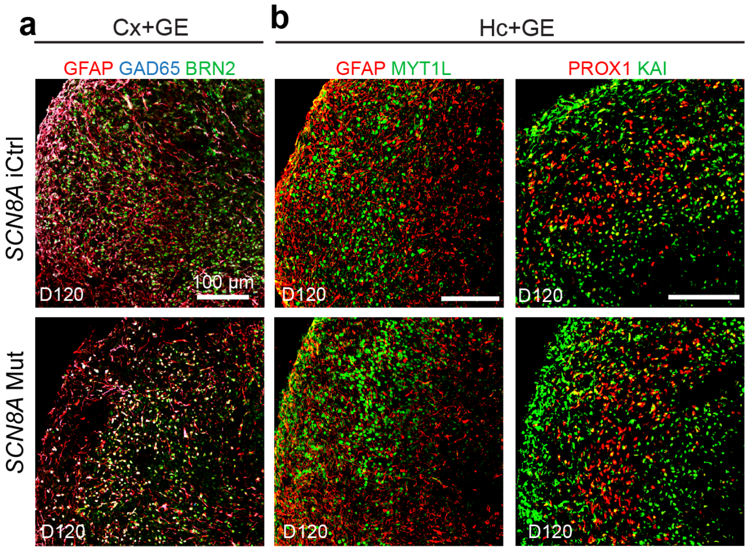


**Extended Data Figure 1- Cx+GE and Hc+GE Immunohistochemistry Demonstrates the Presence of Expected Cell Types: a**, Immunohistochemical analysis of iCtrl and Mut Cx+GE fusion organoids at D120 reveals the presence of glial (GFAP), GABAergic interneuron (GAD65), and a neuronal marker (BRN2). **b**, Immunohistochemical analysis of iCtrl and Mut Hc+GE at D120 reveals the presence of GFAP and the neuronal marker MYTL1, as well as distinct dentate granule (PROX1) and CA3-like (KA1) regions.

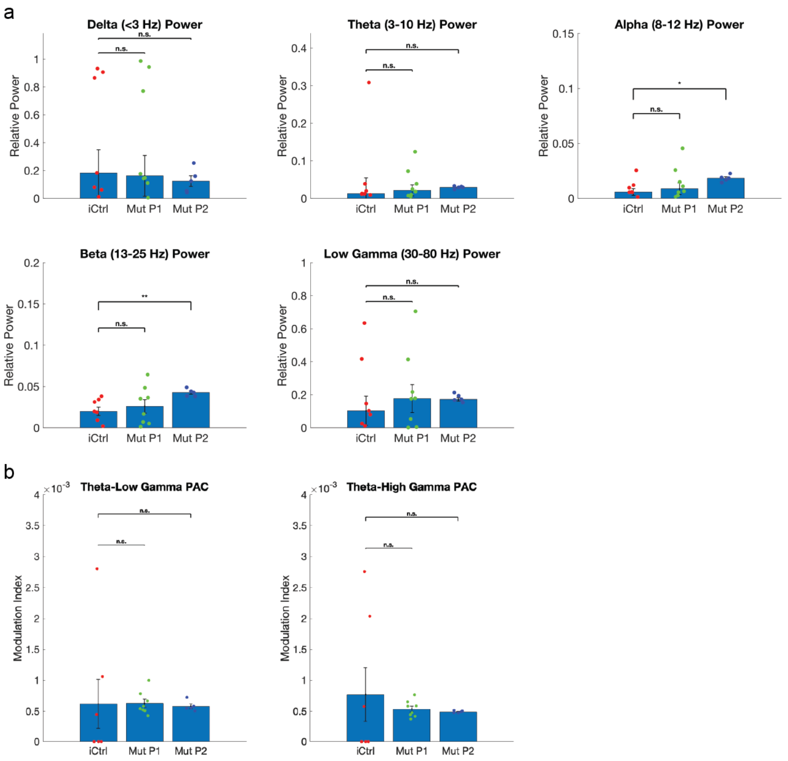


**Extended Data Figure 2- Relative Spectral Power Across Different Frequency Bands and Strength of Theta-Gamma Phase-Amplitude Coupling in Hc+GE Assembloids: a**, Each datapoint represents the median relative spectral power in a given frequency band for a single assembloid, across all 10-second trials available for that assembloid. Spectral power was calculated using Welch's power spectral density estimate. There were no consistent differences in the relative spectral power across canonical frequency bands (delta, theta, alpha, beta, and low gamma) between iCtrl and Mut assembloids. Notably, the only significant differences observed were an increase in relative alpha (8-12 Hz) and beta (13-25 Hz) power in Mut Patient 2 Hc+GE assembloids compared to controls (two-tailed Wilcoxon rank-sum tests). These findings indicate that the disruption of monophasic theta-gamma phase-amplitude coupling in both Mut Patient 1 and Mut Patient 2 Hc+GE assembloids, as shown in Fig. 4 (main text), is likely not attributable to changes in the overall power of oscillatory rhythms in the local field potentials recorded from these assembloids. ns= $p>0.05$, *= $p<0.05$, **= $p<0.01$. **b**, We used the modulation index to measure the strength of coupling between the phase of theta (3-10 Hz) rhythms and the amplitude of both low gamma (30-80 Hz) and high gamma (80-160 Hz) rhythms in the Hc+GE assembloids presented in Figure 4e-g. Notably, there were no significant differences between the iCtrl and Mut assembloids (two-tailed Wilcoxon rank-sum tests).

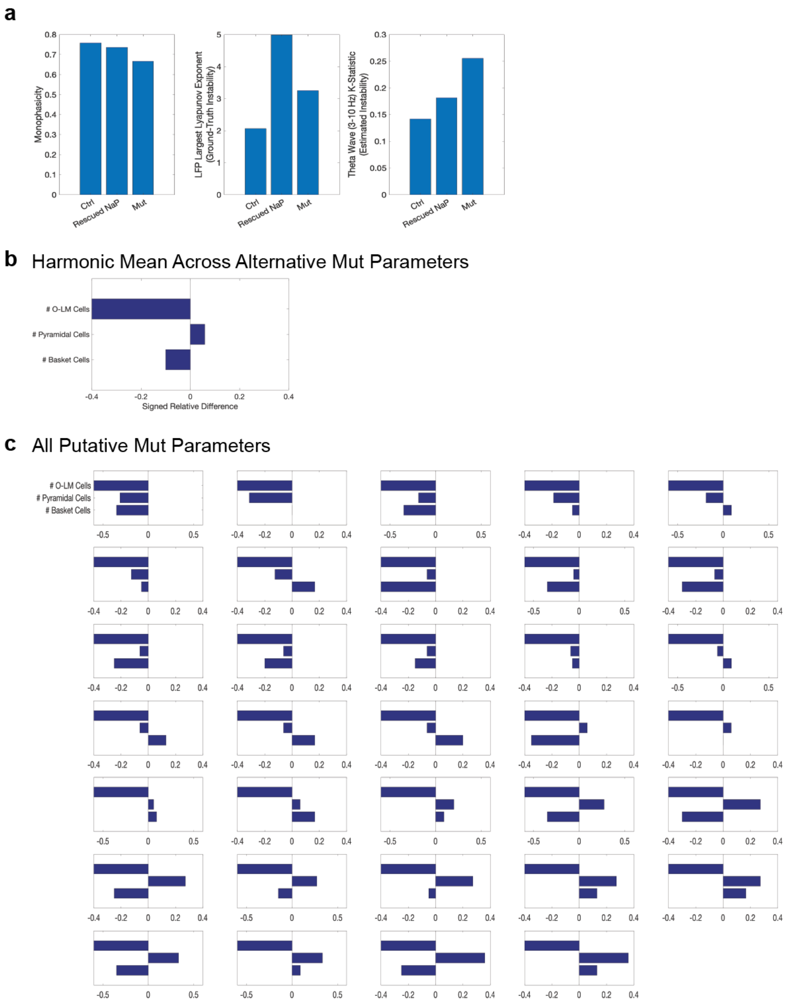


**Extended Data Figure 3- Simulations with Renormalized Persistent Sodium Current and Alternative Mut CA3 Network Composition Parameters: a**, We used our hippocampal model to evaluate, in silico, whether targeted depletion of *p.SCN8A* with NaV1.6 directed anti-sense oligonucleotides or pharmacological blockage of sodium channels might restore hippocampal function, even in the presence of the altered circuit composition identified in the main paper (namely, a loss of O-LM cells and an increase in the number of pyramidal cells). To do so, we simulated our hippocampal circuit using the cell number parameters of our Mut simulation, but fully “rescued” the persistent sodium currents by setting them to the values for our Ctrl simulation. We found that rescuing persistent the persistent sodium currents partially (but not fully) restored the monophasicity of theta-gamma coupling. However, we found inconsistent effects on the stability of theta waves: when using the largest Lyapunov exponent, theta waves were estimated to be *more* unstable in the rescued persistent sodium current simulation than in the full Mut simulation; but, when using the K-statistic of the 0-1 chaos test, rescuing persistent sodium currents partially (but not fully) re-stabilized theta waves. In general, these results suggest that rescuing aberrant sodium conductance in DEE-13 may not be sufficient to fully restore hippocampal function, and point to the potential need for interventions which also target the balance of excitatory and inhibitory cell populations. **b**, We sought to test whether there were any simulated hippocampal circuit compositions, besides the one identified by machine learning (Fig. 6u, main text), that could recapitulate the Mut electrophysiology phenotype (namely, a reduction of monophasic theta-gamma coupling, coupled with a destabilization of both local field potential and single-neuron spiking, with minimal changes to theta or gamma power). To do so, we fixed the persistent sodium current conductances that were identified by machine learning for our Mut simulation (Fig. 6u, main text). We then simulated the hippocampal circuit with 1,280 different combinations of O-LM, basket, and pyramidal cell numbers and identified, among these, which alternative circuit compositions could recreate the Mut Hc+GE assembloid electrodynamic phenotype. As in Fig. 6u, for each putative Mut simulation, we looked at the signed relative difference between the number of each cell type and the corresponding number of cells in the Ctrl simulation. The harmonic mean of these ratios across all putative Mut simulations was largely consistent with the circuit composition identified by machine learning (Fig. 6u), with a marked decrease in O-LM cells and an increase in the number of pyramidal cells. However, there was also a general tendency toward a decrease in the number of basket cells, which is consistent with the IHC results for the Patient 2 Mut Hc+GE assembloids (Fig. 5a, main text). **c**, However, when looking at the circuit composition of each individual putative Mut simulation, we note that the only consistent change across all alternative Mut parameters was a decrease in the number of O-LM cells, with variability both in the number of pyramidal and basket cells, suggesting that a loss of O-LM cells is indeed causal for the aberrant electrophysiological phenotype observed in the Mut Hc+GE assembloids.

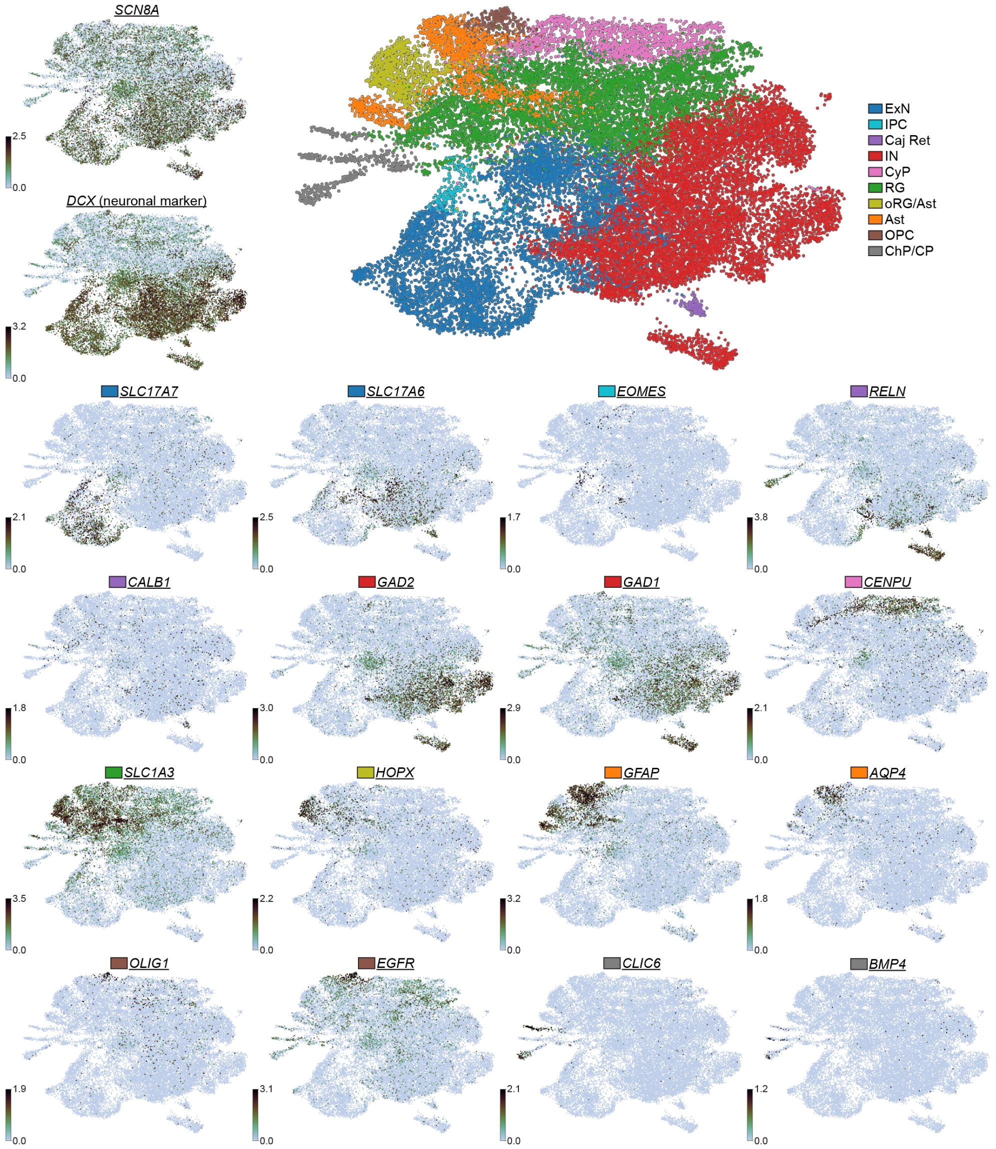


**Extended Data Figure 4-Cortical and Hippocampal Assembloids Contain Diverse Cell Types with Canonical Expression Patterns:** UMAP of all cells from both iCtrl and Mut P1 Cx+GE and Hc+GE assembloids integrated together that demonstrates a rich diversity of cell types and diffuse *SCN8A* expression. The cells were well separated into neuronal (*DCX*-expressing) and non-neuronal cell types. These include *SLC17A6*-/*SLC17A7*-expressing excitatory neurons (ExN), *EOMES*-expressing intermediate progenitors (IPC), *RELN*-/*CALB1*-expressing Cajal-Retzius cells (Caj Ret), *GAD2*-/*GAD1*-expressing inhibitory interneurons (IN), *CENPU*-expressing cycling progenitors (CyP), *SLC1A3*-expressing radial glia

(RG), *HOPX*-expressing outer radial glia that overlapped with *GFAP*-expressing astrocytes in an oRG/Ast co-cluster, as well as astrocytes that were not associated with oRG (labeled Ast) and exhibited more mature expression markers such as *AQP4*, *OLIG1*-/*EGFR*-expressing oligodendrocyte precursor cells (OPC), and cells that expressed choroid plexus-like (*CLIC6*) or cortical plate-like markers.

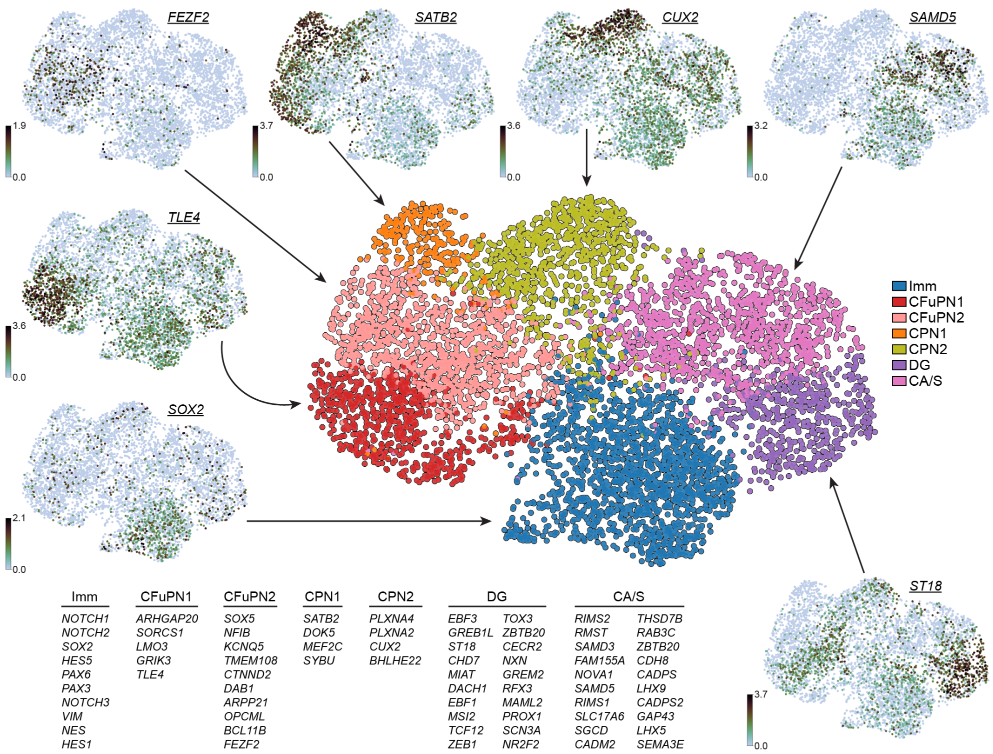


**Extended Data Figure 5-Excitatory Neuron Subtypes in Cortical and Hippocampal Assembloids Show Distinct Expression Patterns:** UMAP of excitatory neuron subtypes from both iCtrl and Mut P1 Cx+GE and Hc+GE assembloids showing well-defined clusters corresponding to 1) immature excitatory neurons (Imm) that express *NOTCH1*, *SOX2*, *HES5*, *VIM*, 2) corticofugal projection neurons (CFuPN1 and CFuPN2) that express *TLE4*, *BCL11B*, *FEZF2*, 3) callosal projection neurons (CPN1 and CPN2) that express *SATB2*, *CUX2*, *BHLHE22* as well as markers of maturity (such as *SYBU*), 4) dentate granule-like cells (DG) that express canonical dentate gyrus and hippocampal markers (*PROX1*, *ST18*, *ZBTB20*), and 5) cornu ammonis/subiculum-like excitatory neurons (CA/S) that express *LHX9*, *THSD7B*, *CADPS*/*CADPS2*, *SLC17A6*. The listed genes are differentially expressed in that cluster ($p - adj < 0.05$) and are listed in order of significance.

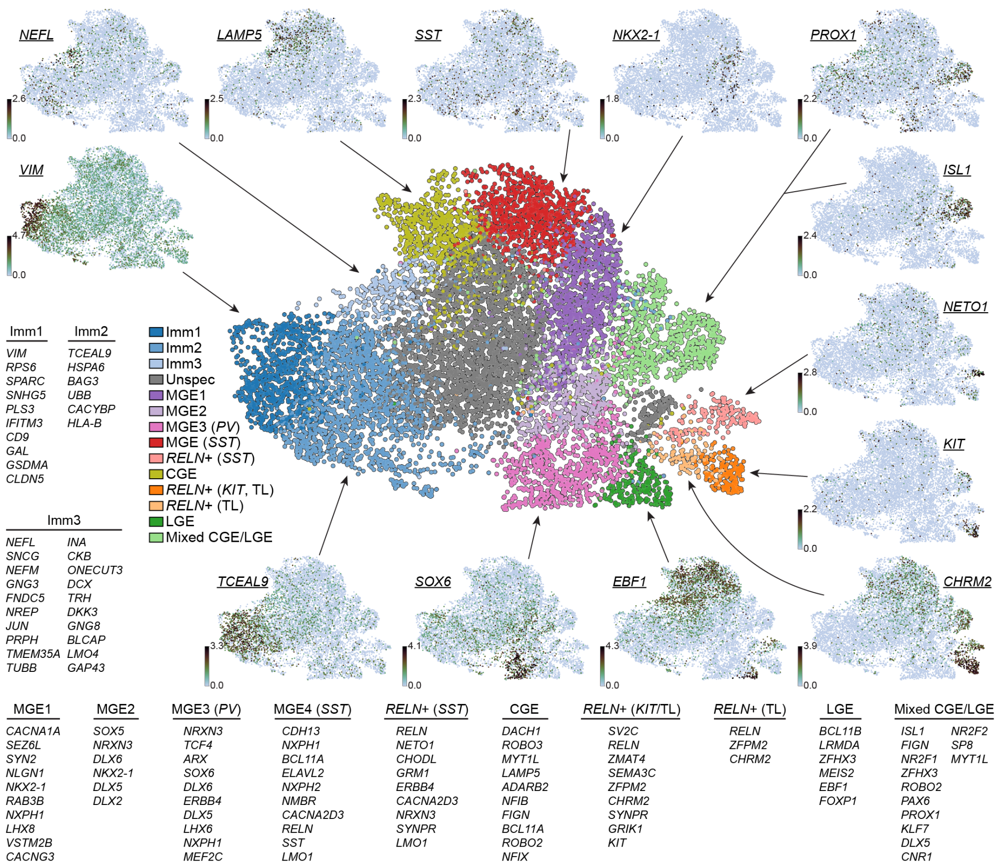


**Extended Data Figure 6-Rich Inhibitory Interneuron Subtype Diversity from Cortical and Hippocampal Assembloids:** UMAP of inhibitory interneuron (IN) subtypes from both iCtrl and Mut P1 Cx+GE and Hc+GE assembloids showing diverse populations. Clusters of immature INs at various stages were present, such as early *VIM*- and *SPARC*-expressing INs (Imm1), later *TCEAL9*- and *BAG3*-expressing INs (Imm2), and even later *NEFL*-, *SEFM*-, and *ONECUT3*-expressing INs (Imm3). A population of unspecified INs (Unspec) was present that lacked well-defined expression markers corresponding to specific known subtypes. Multiple populations of medial ganglionic eminence (MGE)-like INs were present; some cells were less differentiated and expressed *NKX2-1*, *LHX8*, and *DLX6* (MGE1 and MGE2), while others expressed markers of parvalbumin MGE INs (MGE3 (*PV*)), such as *SOX6*, *ERBB4*, *MEF2C*, or somatostatin MGE INs (MGE4 (*SST*)), such as *SST* and *NMBR*. *RELN+* INs that expressed *SST*-type MGE markers, such as *NETO1*, *CHODL*, *GRM1*, were also present (*RELN+* (*SST*)). INs with caudal ganglionic eminence markers such as *LAMP5* and *ADARB2* were also present (CGE). Additionally, *RELN+* INs with CGE-like gene expression profiles that robustly expressed *KIT* and the trilaminar interneuron marker *CHRM2* (*RELN+* (*KIT*/TL)) or *CHRM2* only (*RELN+* (TL)) were present. Lateral ganglionic eminence-like INs were also present that expressed *EBF1*, *LRMDA*, *MEIS2* (LGE). One cluster (Mixed LGE/CGE) that expressed genes canonical to both LGE and CGE INs (*ISL1*, *PAX6* as well as *PROX1*, *NR2F2*) was present. The listed genes are differentially expressed in that cluster ($p - adj < 0.05$) and are listed in order of significance.

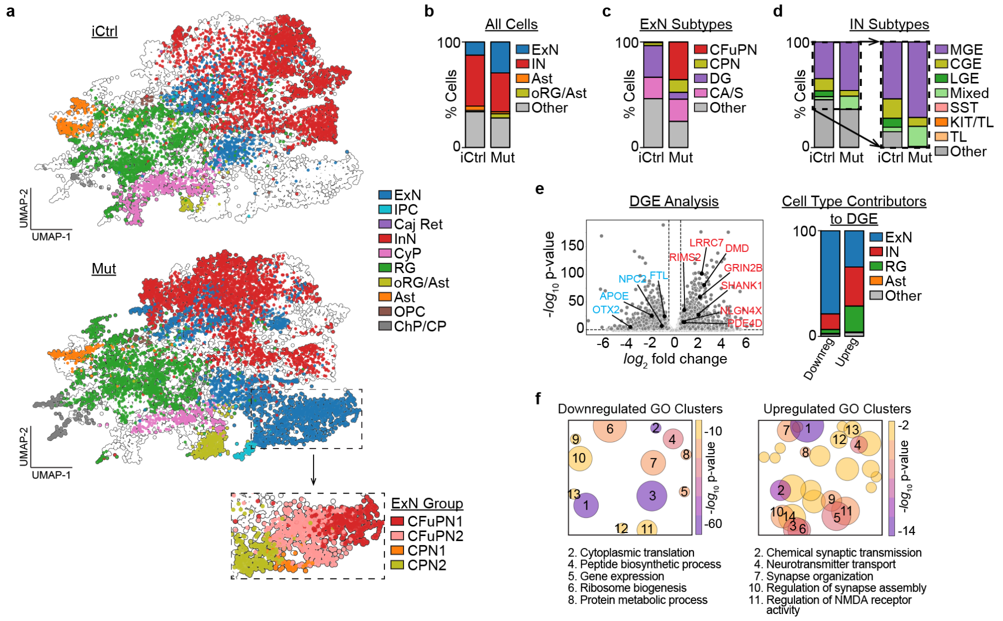


**Extended Data Figure 7- Early Effects of DEE-13 on Circuit Composition and Gene Expression in Hippocampal Assembloids at Day 84 Show Similarities and Differences Versus Day 120 Assembloids: a**, UMAP of cells from iCtrl and Mut P1 Hc+GE showing diverse cell types as well as an increase in the cortex-like excitatory neuron (ExN) clusters, i.e. corticofugal projection neurons (CFuPN) and callosal projection neurons (CPN), in the MutP1 Hc+GE assembloids (see zoomed in box of ExN group). **b**, Mut P1 Hc+GE assembloids show increased numbers of excitatory neurons (ExN) and decreased inhibitory interneurons (IN) compared to iCtrl, similar to the Day 120 Hc+GE assembloids. Likewise, there is a reduction in astrocytes (Ast) and increase in the co-cluster of outer radial glia and astrocytes (oRG/Ast), which was also present in the Day 120 Hc+GE assembloids. **c**, Mut P1 Hc+GE assembloids had an increase in cortex-like CFuPN and CPN ExNs compared to iCtrl and a decrease in the hippocampal dentate granule (DG)-like ExNs, mirroring what was seen at Day 120 although without a substantial reduction in cornu ammonis/subiculum (CA/S)-like ExNs. **d**, There was a mild reduction of medial ganglionic eminence (MGE)-like INs in Mut P1 Hc+GE compared to iCtrl without a large change in the overall caudal or lateral ganglioninc eminence (CGE, LGE respectively)-like INs. No *RELN*+ INs (including SST, KIT/TL, and TL) were present in either iCtrl or Mut P1 Day 84 Hc+GE. **e**, Differential gene analysis volcano plot with a select subset of upregulated genes (red) in Mut P1 assembloids (predominantly in ExNs, INs, and radial glia (RG)) that are involved in glutamatergic pathways and a subset of downregulated genes (blue), predominantly in ExNs, that are associated with epilepsy and intellectual disability when their expression is reduced (from DisGeNET) and are shown in relative proportion compared to those in Fig. 6k and 6l in the main text. **f**, Gene ontology (GO) analysis using all differentially expressed genes and super-clustering of terms demonstrated a upregulation of genes involved in synaptic function in Day 84 hippocampal Mut P1 assembloids compared to iCtrl as well as a broad downregulation of pathways involved in basic cellular viability/housekeeping tasks (e.g. translation, gene expression, metabolism); this was overall similar to the trend observed in Day 120 assembloids although at Day 120 the upregulated synaptic genes were more strongly involved in glutamatergic activity. Larger circles correspond to a higher number of GO terms. **Additional Abbr.** IPC = intermediate progenitor cells, Caj Ret = Caja-Retzius cells, CyP = cycling progenitors, OPC = oligodendrocyte precursor cell, ChP/CP = choroid plexus/cortical plate.

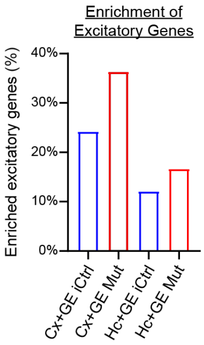


**Extended Figure 8- Increased Excitatory Gene Expression in iCtrl and Mut Cortical Compared to Hippocampal Assembloids:** Enriched expression of excitatory-related genes in iCtrl and Mut Cx+GE and Hc+GE assembloids. There is increased expression in Mut compared to iCtrl for both Cx+GE and Hc+GE, as well as a generalized increased in Cx+GE compared to Hc+GE, consistent with region- and condition-specific differences in excitatory-inhibitory balance.

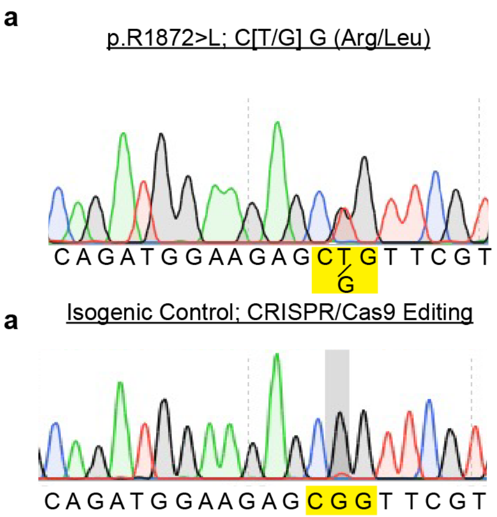


**Extended Figure 9- Sanger Sequencing of p.R1872>L and the CRISPPR-Cas9 Correction:** (a) Sanger sequencing of p.R1872>L demonstrating the T/G point mutation resulting in a Arg/Leu amino acid change (Mut P1). (b) The patient 1 isogenic control (iCtrl) showing successful CRISPR-Cas9 mediated correction of the point mutation.

| Patient Number | Age | Gender | Epileptic side |
|---|---|---|---|
| 1 | 44 | Male | Left |
| 2 | 33 | Female | Right |
| 3 | 25 | Female | Right |
| 4 | 31 | Female | Left |
| 5 | 50 | Male | Right |
| 6 | 39 | Male | Right |
| 7 | 31 | Male | Right |
| 8 | 37 | Female | Left |

**Extended Data Table 1-Temporal Lobe Epilepsy Patient Demographics:** All temporal lobe epilepsy patients whose hippocampal field potentials were analyzed in Fig. 6i-m (main text) received bilateral intracranial depth electrode implantations. However, in each patient, epileptic activity was confined to the hippocampus in one hemisphere, with the hippocampus in the other hemisphere remaining healthy.
