## Supplementary Data for "Modeling Cortical Versus Hippocampal Network Dysfunction in a Human Brain Assembloid Model of Epilepsy and Intellectual Disability"

### 1 Supplementary Videos

Supplementary Videos: Live two-photon imaging of calcium indicator activity within representative assembloids. When indicated, the videos include representative examples of synchronized transients and unsynchronized ("Unsynch") transients. All assembloids were infected with AAV1-GCaMP7f and imaged between 12-16 days later (119-124 DIV) using 2-photon microscopy. The videos are rendered at 10x speed with recording time in seconds indicated on the upper left corner during playback.

Supplementary Video 1: D124 iCtrl Cx+GE- unsynchronized example
Supplementary Video 2: D119 Mut P1 Cx+GE- synchronized example
Supplementary Video 3: D119 Mut P1 Cx+GE- unsynchronized example
Supplementary Video 4: D120 Mut P2 Cx+GE- synchronized example
Supplementary Video 5: D120 Mut P2 Cx+GE- unsynchronized example
Supplementary Video 6: D123 iCtrl Hc+GE- synchronized example
Supplementary Video 7: D120 iCtrl Hc+GE- unsynchronized example
Supplementary Video 8: D120 Mut P1 Hc+GE- synchronized example
Supplementary Video 9: D120 Mut P1 Hc+GE- unsynchronized example
Supplementary Video 10: D120 Mut P2 Hc+GE- synchronized example
Supplementary Video 11: D122 Mut P2 Hc+GE- unsynchronized example
